# Salt supersaturation as accelerator of influenza A virus inactivation in 1-μl droplets

**DOI:** 10.1101/2023.12.21.572782

**Authors:** Aline Schaub, Beiping Luo, Shannon C. David, Irina Glas, Liviana K. Klein, Laura Costa, Céline Terrettaz, Nir Bluvshtein, Ghislain Motos, Kalliopi Violaki, Marie Pohl, Walter Hugentobler, Athanasios Nenes, Silke Stertz, Ulrich K. Krieger, Thomas Peter, Tamar Kohn

## Abstract

Influenza A virus (IAV) spreads through exhaled aerosol particles and larger droplets. Estimating the stability of IAV is challenging and depends on factors such as the respiratory matrix and drying kinetics. Here, we combine kinetic experiments on millimeter-sized saline droplets with a biophysical aerosol model to quantify the impact of NaCl on IAV stability. We show that IAV inactivation is determined by NaCl concentration, which increases during water evaporation and then decreases again when efflorescence occurs. When drying in air with relative humidity RH = 30%, inactivation follows an inverted sigmoidal curve, with inactivation occurring most rapidly when the NaCl concentration exceeds 20 molal immediately prior to efflorescence. Efflorescence reduces the NaCl molality to saturated conditions, resulting in a significantly reduced inactivation rate. We demonstrate that the inactivation rate *k* depends exponentially on NaCl molality, and after the solution reaches equilibrium, the inactivation proceeds at a first-order rate. Introducing sucrose, an organic co-solute, attenuates IAV inactivation via two mechanisms, firstly by decreasing the NaCl molality during the drying phase, and secondly by a protective effect against the NaCl-induced inactivation. For both pure saline and sucrose-containing droplets, our biophysical model ResAM accurately simulates the inactivation when NaCl molality is used as the only inactivating factor. This study highlights the role of NaCl molality in IAV inactivation and provides a mechanistic basis for the observed inactivation rates.

SYNOPSIS: This work quantifies the dependence of influenza A virus stability on salt molality in drying droplets and furthers the understanding of airborne virus transmission.SYNOPSIS: This work quantifies the dependence of influenza A virus stability on salt molality in drying droplets and furthers the understanding of airborne virus transmission.

## INTRODUCTION

Respiratory viruses such as influenza A virus (IAV) can be transmitted by expiratory aerosol particles and droplets emitted by an infected individual.^1–5^ Small particles (up a few μm) can stay airborne for minutes to hours^6^ and travel dozens of meters before encountering a new host,^7^ while larger droplets (> 100 μm) can travel only a short distance (1-2 m) and settle within seconds.^7^ For both aerosol particles and droplets, IAV can only be transmitted if the viruses remain infectious outside the host. Understanding the environmental parameters that modulate IAV infectivity in expiratory particles and droplets is therefore crucial to tackle seasonal flu epidemics as well as worldwide pandemics.

When aerosol particles and droplets are exhaled from the respiratory tract, they leave an environment with a relative humidity (RH) close to 100% and body temperature and instantaneously enter a cooler, dryer environment. In typical indoor environments in temperate climates, RH can be as low as 30% during winter, whereas it is around 50% in summer.^8^ The RH,^9–19^ the temperature,^20–22^ and the initial size and composition of the expiratory particle^23–25^ dictate the droplet drying rate and hence the concentration of solutes in the droplets. If the RH is below the efflorescence relative humidity (ERH) of the salts, the salts will effloresce. Moreover, gas exchange with the environment takes place,^25^ and liquid-liquid phase separation may occur.^26^ These processes modulate the salinity as well as the pH within the particle. We previously found that the pH in small aerosol particles rapidly decreases to around 4 as a result of gas exchange with the surrounding air.^27^ This low pH was suggested to be a primary driver of IAV inactivation in aerosol particles in indoor environments.^27,28^ In contrast, larger droplets first undergo alkalization^29^ up to pH 10 followed by a much slower acidification on the scale of hours,^25^ suggesting that IAV inactivation observed in droplets is likely driven by another factor.

Previous studies have suggested an important contribution of salt to virus inactivation in deposited droplets. Specifically, the stability of IAV,^30^ bacteriophage MS2,^18^ and SARS-CoV-2^20^ was found to be low at intermediate RH (50-65%), just above ERH of NaCl (41-51%),^31^ where the salinity in the droplets is expected to be maximal. In search for a mechanistic explanation for this observation, Lin and Marr^18^ suggested that the cumulative dose (NaCl concentration multiplied by exposure time) of NaCl over a 1-h drying period is the main determinant of virus inactivation in droplets and aerosol particles for bacteriophages MS2 and Φ6. Similarly, Niazi et al.^32,33^ showed that rapid evaporation followed by immediate efflorescence does not inactivate IAV and human rhinovirus-16 in airborne aerosol particles, suggesting that a short exposure to high salinity is insufficient to lead to efficient inactivation. An important role of evaporation dynamics on IAV inactivation in droplets was also proposed by French et al.,^23^ who observed faster inactivation during the initial evaporation step, while the particle is still liquid, and a slower decay after efflorescence. Likewise, a near-instantaneous 50% loss in SARS-CoV-2 infectivity was observed when efflorescence occurred, followed by a slower inactivation rate.^34,35^

Besides salt efflorescence, IAV stability is also dependent on the presence of organic co-solutes in the matrix, which are typically reported to exert protective effects,^30,36^ though the underlying mechanisms are not understood. Nevertheless, some organic compounds, such as the saliva constituents lysozyme and lactoferrin, are viricidal and can enhance viral inactivation.^37,38^ Finally, Rockey et al.^39^ observed a lower IAV stability in saliva droplets than in respiratory mucus droplets, whereby the difference could not be explained by the salt content nor the drying kinetics, but was instead attributed to the morphology of the dried droplets. Overall, the role of salinity, efflorescence and organic co-solutes in IAV inactivation is thus not fully understood, and it remains difficult to predict inactivation in saline droplets, in particular in the presence of organics.

The goal of this study was therefore to provide a mechanistic understanding of the role of salt in IAV inactivation in deposited, drying droplets, to explain observed differences in inactivation rates at different RH and solution conditions. To this end, we aimed to establish a predictive relationship between the NaCl concentration in drying droplets and IAV stability, in the presence and absence of organics.

## MATERIAL AND METHODS

### Virus propagation, purification and enumeration

Influenza A virus (IAV) strain A/WSN/33 (H1N1 subtype) was propagated in Madin-Darby Canine Kidney (MDCK) cells and purified by ultracentrifugation as previously described^27^ and detailed in the Supporting Information (SI). Virus infectivity was quantified by plaque assay. Briefly, a 12-well plate containing MDCK monolayers was washed with phosphate-buffered saline (PBS; ThermoFisher, 18912014) and infected with IAV samples. The plates were incubated for 72 h (37°C, 5% CO_2_) and were then fixed and stained as described in the SI to enumerate and determine the virus titer in plaque forming units (PFU)/ml. A negative control without added virus was always performed on an additional plate. The assay limit of quantification (LoQ) was 10 PFU/ml. The total virus concentration was quantified as genome copies (GC)/ml by Reverse Transcription quantitative Polymerase Chain Reaction (RT-qPCR) as detailed in the SI. All RT-qPCR procedures followed MIQE guidelines^40^ (see **Table S1**).

### Virus inactivation experiments in droplets

While working with real respiratory matrices is the gold standard, we here aimed to ensure tight control on salinity and organic content in our experiments. We therefore chose to work with a model matrix consisting of NaCl at physiological concentration, and sucrose as an organic substance that can modulate the occurrence of efflorescence. We monitored the inactivation of influenza virus strain A/WSN/33 in 1- and 2-μl drying droplets containing NaCl and sucrose at different ratios. A spherical 1-μl droplet has a radius of 620 μm and thus represents the upper end of the particle size distribution produced while coughing or speaking.^41,42^ This volume also allows to contain a sufficient number of viruses to monitor several orders of magnitude (log_10_) of inactivation.

Experiments were conducted in solution with an initial composition of either 8 g/l (i.e., molality *b* = 0.14 mol/(kg H_2_O) = 0.14 molal) NaCl solution (ThermoFisher Scientific, 207790010) or 5.8 g/l (0.14 molal) LiCl solution (Acros Organics, 193370010). This salt concentration corresponds to the NaCl concentration in simulated lung fluid, that was used previously to assess IAV inactivation in a realistic matrix,^27^ falls within the range of salt concentrations measured in sputum (5.9–8.9 g/l),^43^ and is also really close to the NaCl concentration in PBS (8.12 g/l). Additionally, experiments were conducted in solutions containing 8 g/l NaCl solution supplemented with either 8 g/l (0.023 molal) or 64 g/l (0.187 molal) of sucrose as a model organic (Fisher BioReagents, BP220-1). These sucrose concentrations were chosen based on droplet efflorescence analysis (**Figure S1**), which showed that the lower sucrose concentration allowed for NaCl efflorescence, whereas no efflorescence was observed (by videography or eye) at the higher one. The solutions were all prepared using milli-Q water. Purified virus stock in PBS was spiked into 100 μl of the matrix in a 1.5-ml plastic tube (Sarstedt, 3080521), to achieve a starting concentration in the experimental solution of either 10^7^ PFU/ml (“normal titer”) or 10^9^ PFU/ml (“high titer”). The added volume of PBS (1%) does not influence the final concentration of NaCl, as the NaCl content of PBS corresponds closely to the initial NaCl concentration. The tube was vortexed and kept in a cold tube rack until use.

Inactivation experiments were performed in an environmental chamber (Electro-Tech Systems, 5532) with controlled relative humidity and temperature. Prior to the experiment, the chamber was set at the targeted RH and room temperature (25°C ± 2) and left to equilibrate. Trace gases potentially present in the chamber air (e.g. acids such as HNO_3_), which are absorbed by the droplets, have no influence on the inactivation process, as the large volume of the droplets leads to a high dilution. Once equilibrated, 1- or 2-μl droplets were deposited in the wells of a 96-well plate with a non-binding surface (Greiner Bio-One, 655901), one droplet per well. Droplets were then left to dry at 30% or 65% RH (unless mentioned otherwise), and were periodically collected (see below) over the course of up to 180 min. For each time point in an experiment, three droplets were collected. The three droplets that were deposited last were sampled immediately to act as the time = 0 sample. The deposition time (∼10 s per droplet) was recorded and accounted for when sampling the time points. In each experiment, a control was performed in the experimental solution in bulk.

To sample a droplet, 300 μl of PBSi (PBS for infection; PBS supplemented with 1% P/S, 0.02 mM Mg^2+^, 0.01 mM Ca^2+^, and 0.3% bovine serum albumin (BSA, Sigma-Aldrich A1595), with a final pH of ∼7.3) were added to the corresponding well. Each droplet was then resuspended by scratching the bottom of the well with a 200-μl pipette for 5 s, followed by up and down pipetting for 5 times, followed by a second round of scratching and pipetting. Then, each sample was aliquoted in 2x 150 μl and frozen at -20°C until further quantification by culturing and RT-qPCR. The recovering and freezing processes do not inactivate the virus further. In each experiment, a control was included by sampling 1 μl of the experimental solution directly from the plastic tube at the beginning and at the end of the experiment. The sample was immediately diluted in 300 μl of PBSi. The tube was kept in the environmental chamber during the whole experiment to ensure similar conditions as for the droplets.

To account for physical loss of the virus due to attachment to the wells, recovery of the total (infectious plus inactivated) virus from the plate was measured by RT-qPCR. The measured inactivation was then corrected for virus recovery to determine the actual inactivation.

### Virus inactivation experiments in bulk

Inactivation in NaCl bulk solutions were performed in triplicate at room temperature, at three different NaCl concentrations (for subsaturated 0.14 molal and 2.8 molal solutions, and for the saturated 6.1 molal solution). A purified virus stock in PBS was spiked in 1 ml of each matrix in a 1.5-ml plastic tube to achieve a starting concentration in the experimental solution of 10^7^ PFU/ml. The tubes were vortexed and a 5-μl sample was taken immediately and diluted in 495 μl of PBSi. Samples were taken every 2 h for 10 h, and a final time point was taken after 24 h. All the samples were frozen at -20°C immediately after sampling until quantification.

### Determination of inactivation rate coefficient

Inactivation rate coefficients of each experimental condition were determined from least square fits to the log-linear portion of the inactivation curve (that is, for droplets, the inactivation occurring in equilibrated droplets, i.e. ≥ 40 min after deposition), assuming first order kinetics,

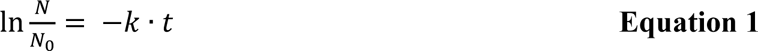

Here, *N* is the number of infectious viruses in a droplet at time *t*, *N*_0_ is the initial number of viruses in the droplet and *k* is the inactivation rate coefficient. All replicate experiments of a given experimental condition were pooled. 99%-inactivation times (*t*_99_) were determined based on *k*:

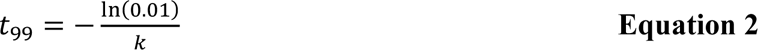

Rate coefficients and associated 95% confidence intervals were determined using GraphPad Prism v.10.0.2.

### Determination of evaporating droplet radius

The evaporation of the droplets was filmed by a camera (Sony IMX477R) connected to a Raspberry Pi 4 computer (Model B Rev. 1.4) and a power bank (Varta). The camera was positioned 15 cm above the well plate and took a picture every 16 s. The covered area of each droplet was calculated from the pictures using the software ImageJ (version 1.53t). Briefly, the stack of images was cropped around the droplet of interest and was converted to 8-bit type. Then, an auto-threshold using the “triangle” method was applied across the image stack. A particle analysis was performed on the droplet using default settings, yielding the droplet area of each droplet in pixels. This enables the calculation of relative radii of the (circular) area across all images in the stack. Here, radius (*r*) refers the base of the cap-shaped droplet after being deposited in one of the non-binding wells as measured by video recording paired with ImageJ analysis and normalized by the initial base radius of the droplet (*r*_0_). Obvious outliers within each time course (abnormally low or high values as determined by visual inspection) were excluded. In addition, to improve readability of the figures, only 1 out of 2 data points are shown in the evaporation phase, and 1 out of 5 data points are shown in the equilibrium phase (tail). The analysis of the droplet base radius had a small relative standard deviation of 2.4% on average for a single droplet, and of 12% across different droplets (**Figure S2**).

### Biophysical modeling

We used the Respiratory Aerosol Model ResAM to predict NaCl and sucrose concentrations in the evaporating droplets and to simulate the inactivation kinetics of IAV. ResAM was developed as a spherical shell diffusion model to determine viral inactivation in spherical aerosol particles and droplets, which we generalized here to take account for the nonspherical droplet-in-well symmetry. ResAM is initialized with the starting composition and size of the droplets, as well as the relative humidity, temperature and ventilation in the experimental chamber, and assuming the viruses to be initially homogeneously distributed in the droplet volume. From this initial state, the model simulates the loss of water molecules by gas phase diffusion, the buildup of internal NaCl concentration gradients in the liquid phase, and derives the inactivation rates as function of NaCl molality, taking into account the internal concentration gradients. ResAM is fully described by Luo et al.^27^ and the adaptation to non-spherical droplet geometry is described in detail in the SI, where the uncertainties and model limitations are also discussed.

## RESULTS

### IAV inactivation kinetics in efflorescing NaCl droplets

Inactivation kinetics of IAV in 1- and 2-μl aqueous NaCl droplets exposed to an RH of 30% were measured over the course of 60 min, to compare the kinetics in two different droplet sizes. Simultaneously, the droplet size was monitored as a function of time by videography (**Figure 1**). Initially, the droplets were composed of 8 g/l of NaCl, and 1% (by volume of infectious virus in phosphate buffer saline (PBS), reaching a virus titer of 10^7^ PFU/ml. After deposition, the droplets shrank slowly due to water loss until the salt visibly effloresced after approximately 20 min in the 1- μl droplets and after 36 min in the 2-μl droplets (**Figure 1A+C**). After efflorescence, the appearance of the droplets, as seen by eye and videography, did not change until the end of the experiment.

**Figure 1:**
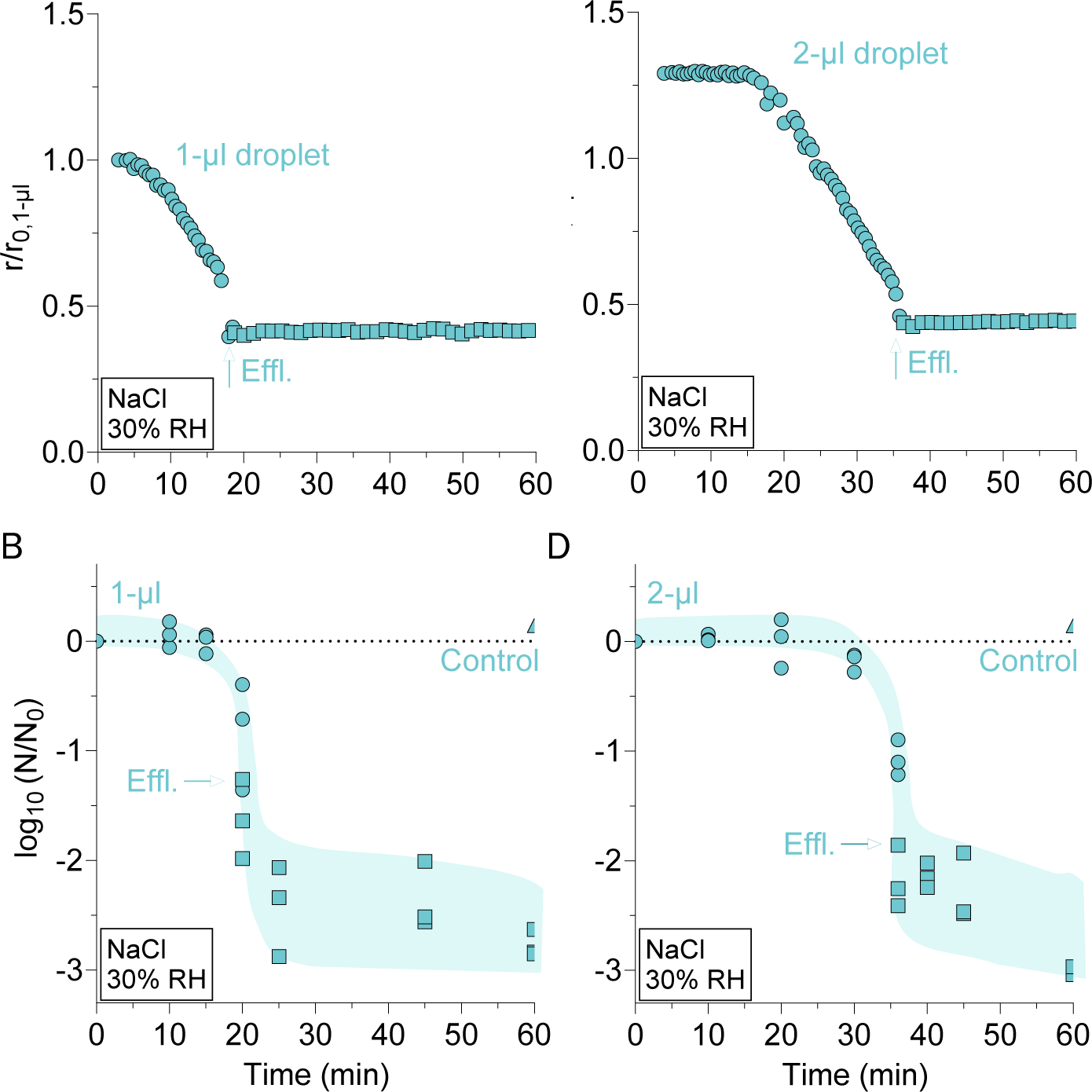
Water loss of highly diluted 1-μl and 2-μl aqueous NaCl droplets ([NaCl]_initial_ = 8 g/l) exposed to 30% RH and concomitant inactivation of IAV in these droplets. Circles represent liquid droplets, squares represent effloresced droplets, and triangles represent control experiments conducted in bulk solution. Efflorescence of NaCl is marked by a green arrow. (A,C) Representative examples of droplet radius shrinkage for one individual droplet with an initial volume of 1 μl (A) and 2 μl (C). Droplet base radius (*r*) was determined by video recording and was normalized by the initial radius of the 1-μl droplet (*r*_0,1-μl_). It first decreases due to evaporation and then remains constant after efflorescence. (B,D) Inactivation of IAV in 1-μl (B) and 2-μl (D) droplets. Each data point represents one individual droplet. Shadings serve to guide the eye and to highlight the small data scatter before, and the large scatter after efflorescence caused by small differences in the stochastic nucleation process triggering efflorescence.

The inactivation curve of IAV in both droplet sizes initially followed the evaporation kinetics of the droplets and exhibited an inverted sigmoidal shape (**Figure 1B+D**): during the initial phase of exposure to the controlled RH in the gas phase, the volume loss of the droplets is greatest, as there is a large humidity gradient between the air directly above the liquid surfaces and the prescribed RH in the environmental chamber. However, during the initial 5-15 min, the camera barely detected any change in droplet size (**Figure 1A+C**). This is because the droplet flattens out, its contact angle with the surface decreases and its volume rapidly decreases, while the covered base area remains constant. This cannot be detected from the elevated angle of the camera. Only after the contact angle reached a critical small size, also the covered base area contracts until equilibrium between the liquid and gas phase in the chamber is reached (i.e., until the water activity in the solution *a_w_* = RH).

Despite the continuous decrease in volume and the increasing salt concentration, IAV remained infectious during the first 15-25 min. This was followed by a short phase of about 5 min during which IAV infectivity plummeted by more than 2-log_10_. After efflorescence, the inactivation kinetics slowed significantly and showed a constant inactivation rate for the remainder of the experiment, resulting in less than 2-log_10_ inactivation in the 40 min after efflorescence. In 2-μl droplets, inactivation kinetics were similar to 1-μl, although the initial slow inactivation phase was prolonged to approximately 30 min, due to the smaller relative volume change rate and, hence, the later efflorescence of the larger droplets. Control experiments conducted in bulk solutions showed only minor inactivation during the course of the experiment, suggesting that the observed inactivation can be attributed to the droplet environment.

Two mechanisms may explain the rapid decease in virus titer. Either, this may be caused by the formation of salt crystals during efflorescence, as suggested in the context of influenza vaccine microneedles^44^ and salt-coated filters on surgical masks.^45^ Or, alternatively, inactivation may be driven by IAV exposure to high salinity just before efflorescence. The dissolved NaCl concentration increases as the droplet evaporates and massive supersaturations (> 20 molal, see below) are reached prior to efflorescence, exposing the virus to extreme salinity, before it decreases again to the saturation level (6.1 molal) due to efflorescence. The latter mechanism, first hypothesized by Yang et al.,^30^ is proven by our finding that substantial inactivation occurs before efflorescence.

Furthermore, **Figure 1B+D** suggests that the data scatter is small before efflorescence, but large afterwards. This observation is also consistent with salinity being the main driver of inactivation: the fastest inactivation occurs under the most saline conditions just before efflorescence, and because efflorescence is a stochastic nucleation process, different droplets effloresce at slightly different times. As a consequence, the viruses in these different droplets are exposed to differently high salt concentrations, resulting in different titers in the moment of efflorescence and beyond.

### IAV inactivation kinetics in non-efflorescing NaCl droplets

To confirm that the main driver of inactivation is high salinity, we tested IAV inactivation under experimental conditions of increasing salt molality but in the absence of efflorescence, combined with droplet size measurements (**Figure 2**). First, experiments were conducted in the same NaCl matrix as in **Figure 1**, but at RH = 65% >> ERH (**Figure 2A-B**). Second, inactivation was measured at RH = 30% (same as in **Figure 1**), but in a matrix containing LiCl instead of NaCl (**Figure 2C-D**). LiCl has an ERH of 1-4%,^31^ and therefore remains deliquesced at 30% RH. As a reference, experiments in NaCl droplets at 30% RH were also included. In this set of inactivation experiments (but not droplet size measurements), the initial virus titer was increased by 2-log_10_ compared to the experiments shown in **Figure 1**, in order to increase the detection range. The initial LiCl molal concentration (0.14 m) was identical to the NaCl molal concentration.

**Figure 2:**
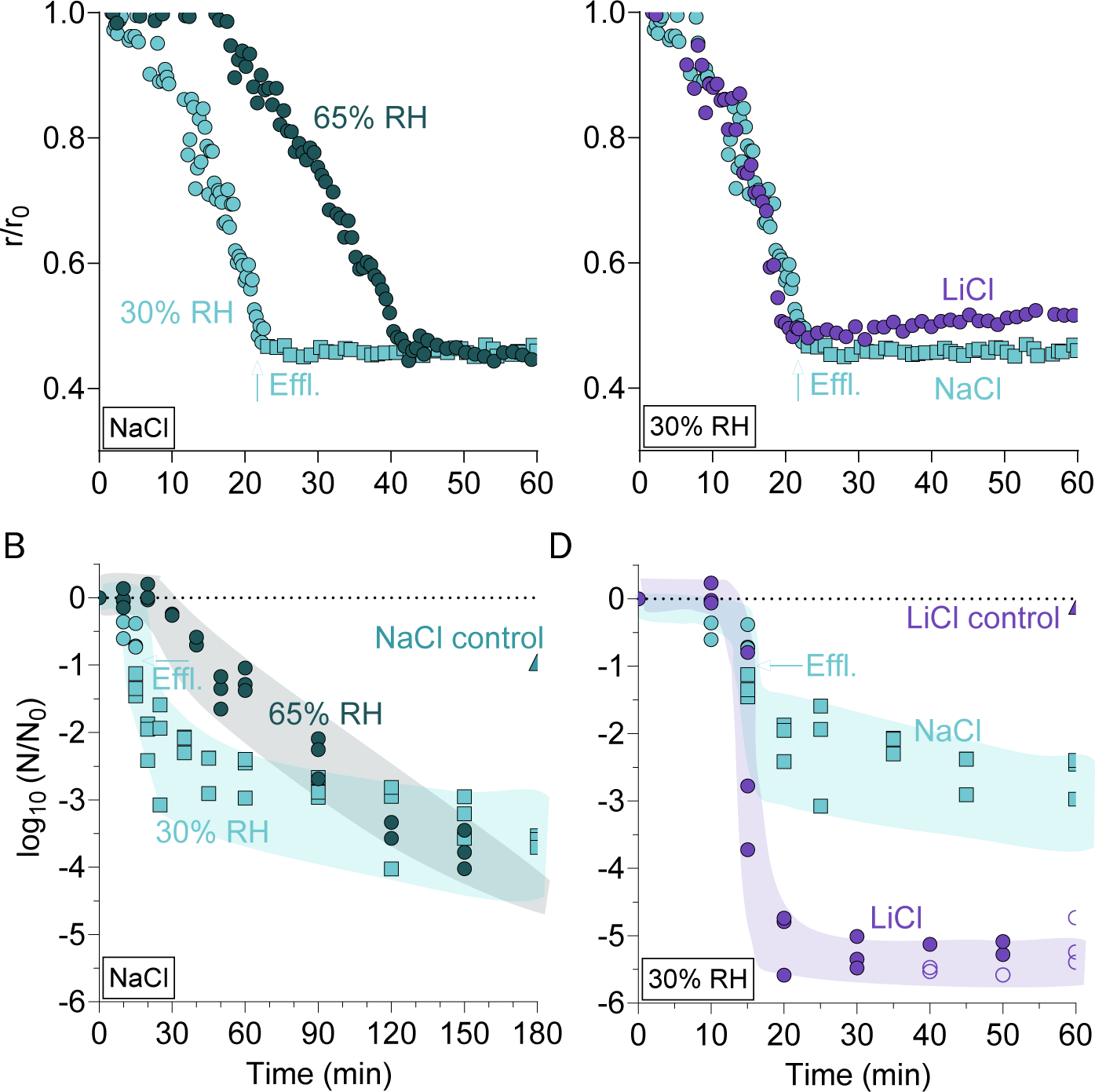
IAV inactivation in 1-μl NaCl droplets at 30% RH (light green) and 65% RH (dark green) for 180 min and in 1-μl LiCl droplets at 30% RH (purple) for 60 min. Circles represent liquid droplets, squares represent effloresced droplets, and triangles represent control experiments conducted in bulk solution. Efflorescence of NaCl (when present) is marked by a green arrow. (A) Representative examples of NaCl droplet radius shrinkage at 30 and 65% RH due to water evaporation until efflorescence (at 30% RH) or liquid- phase equilibrium with air (at 65% RH). Radius (r) was determined by video recording and was normalized by dividing by the initial radius of each droplet (r_0_). (B) Inactivation of IAV in 1-μl NaCl droplets ([NaCl]_initial_ = 8 g/l) at 30% RH and 65% RH. Note the longer time scale shown in this panel, as inactivation progressed beyond 60 min. (C) Representative examples of NaCl and LiCl droplet radius shrinkage at 30% RH due to water evaporation until efflorescence (NaCl) or liquid-phase equilibrium with air (LiCl). (D) Inactivation of IAV in 1-μl LiCl droplets ([LiCl]_initial_ = 5.8 g/l) at 30% RH. Empty symbols indicate data below the LoQ. The control for NaCl was taken after 180 min and is shown in panel B. In panels B + D, each data point represents one individual droplet. Shadings serve to guide the eye. Note that the droplet size measurements (panels A and C) were performed at a lower IAV titer and hence efflorescence occurred about 5 min later than in inactivation experiments (see text).

The inactivation in NaCl droplets at 30% RH followed kinetics comparable to those shown in **Figure 1**, though efflorescence occurred approximately 5 min earlier than in experiments with a lower virus titer (15 min after deposition instead of 20 min, respectively). This suggests that the presence of concentrated virus can trigger efflorescence (**Figure S3**). The first-order inactivation rate coefficient (*k*) after efflorescence (time after deposition ≥ 40 min) was 1.8 (± 0.4) × 10^-2^ min^-1^, corresponding to a time to reach 99% inactivation *t*_99_= 260 min (**Table S2**). As explained for **Figure 1C-D**, the large data scatter after efflorescence is to be expected owing to the stochastic nature of the heterogeneous nucleation process.

The viral infectivity in NaCl droplets at 65% RH remained stable during 20 min after deposition, consistent with initial conditions of high dilution. Subsequently, it exhibited a first-order inactivation with a rate coefficient *k* = 6.5 (± 0.4) × 10^-2^ min^-1^, corresponding to *t*_99_ ≍ 71 min (**Table S2**). After 120 min, IAV inactivation in droplets dried at 30% and 65% RH reached similar levels. In contrast, LiCl droplets at 30% RH exhibited a rapid drop in IAV titer between 10 and 20 min, similar to the time scale in NaCl droplets at the same RH. However, the titer dropped more extensively, and inactivation reached > 5-log_10_ after 30 min. Thereafter, a reduction in the inactivation rate was observed until 60 min, after which the viral titer was no longer quantifiable. Note that decelerating inactivation curves are often observed in experiments leading to a high titer reduction, and may be explained by the presence of viral aggregates.^46,47^ As expected, no efflorescence occurred in the LiCl droplets. Taken together, these experiments indicate that the high salt molality and not the formation of salt crystals causes the inactivation.

### IAV inactivation kinetics in NaCl/sucrose droplets

To assess the inactivation of IAV in the presence of organics, sucrose was added to the NaCl solution. Organics have previously been observed to be protective for IAV in bulk solutions,^27^ deposited droplets,^30^ or airborne aerosol particles.^36^ In these studies, however, the protective matrix was a complex mixture of organics. Here, we chose to use a simple NaCl/sucrose/water system, to be able to determine the concentration of the solutes along the experiment using a biophysical aerosol model (see below). NaCl:sucrose mass ratio of 1:1 and 1:8 were selected (corresponding to 8 and 64 g/l of sucrose, respectively), providing conditions where crystallization occurred (1:1) or was suppressed (1:8) when RH = 30%. The inactivation kinetics of IAV in these two NaCl/sucrose matrices were compared with the inactivation in pure NaCl droplets, here denoted NaCl:sucrose 1:0.

After deposition, the 1:1 NaCl:sucrose droplets shrank until efflorescence occurred after 18 min (**Figure 3A**). The inactivation kinetics exhibited an inverted sigmoidal shape, with a fast decay close to the onset of efflorescence, followed by a shallower tail (**Figure 3B**). In the tail, *k* stayed small and constant (slow first-order inactivation) until conclusion of the experiment. The inactivation kinetics was then quite similar to the one observed in 1:0 droplets (though efflorescence in the 1:1 droplets occurred 3 min later and was associated with a larger droplet radius, a smaller drop in titer and slightly slower first-order inactivation rate coefficient in the tail with 1.8 (± 0.4) × 10^-2^ min^-1^ vs. 7.6 (± 3.3) × 10^-3^ min^-1^ for 1:0 and 1:1 ratios, respectively (**Table S2**)). The 1:8 NaCl:sucrose droplets initially shrank at the same rate as the 1:1 droplets but then equilibrated as liquid droplets with the largest relative radius (**Figure 3A**). The inactivation kinetics was slower than in the 1:0 and 1:1 droplets, reaching less than 1-log_10_ inactivation after 180 min, and an inactivation rate coefficient in the tail of 6.4 (± 1.7) × 10^-3^ min^-1^ (**Table S2**). In both sucrose-containing droplets, the viruses were thus partially protected from NaCl-mediated inactivation, whereby the protection was greater at a higher sucrose content.

**Figure 3:**
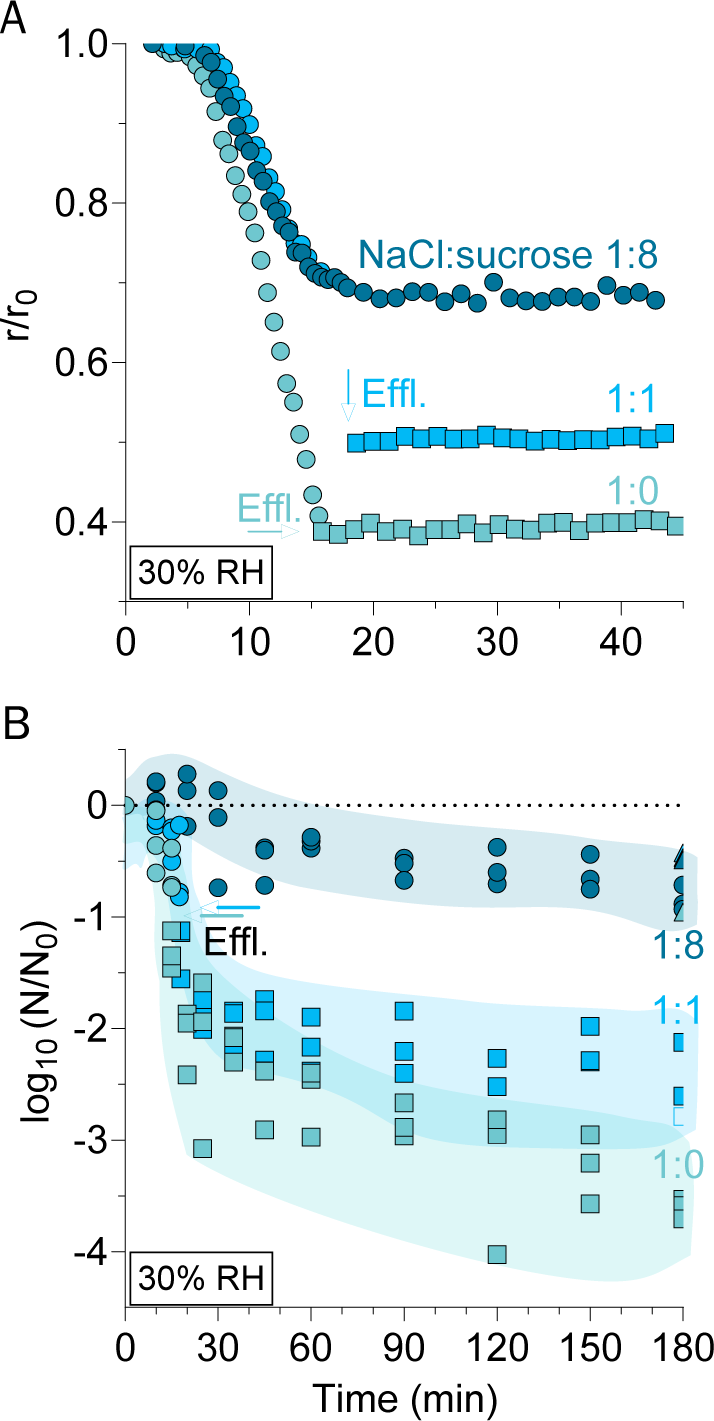
IAV inactivation in 1-μl droplets at 30% RH in various NaCl:sucrose mass ratios. Circles represent liquid droplets, squares represent effloresced droplets, and triangles represent control experiments conducted in bulk solution. Efflorescence, when observed, is indicated by arrows. (A) Representative examples of droplet shrinkage due to water evaporation until efflorescence or liquid-phase equilibrium with chamber RH occurs. Base radius (*r*) was determined by video recording and was normalized to the initial radius of each droplet (*r*_0_). (B) Inactivation of IAV in 1-μl droplets at three different NaCl:sucrose ratios: 1:0 (light green; [NaCl]_initial_ = 8 g/l, [sucrose] = 0 g/l), 1:1 (light blue; [NaCl]_initial_ = 8 g/l, [sucrose]_initial_ = 8 g/l) and 1:8 (dark blue; [NaCl]_initial_ = 8 g/l, [sucrose]_initial_ = 64 g/l). Each data point represents one individual droplet. Empty symbols indicate data below the LoQ. Shadings serve to guide the eye. Note the different time scales in the two panels for improved readability.

### Biophysical model of virus inactivation in drying NaCl/sucrose droplets

The biophysical model ResAM (Respiratory Aerosol Model) was used to simulate the water activity and composition within the droplet as a function of time. A full description of ResAM is available in Luo et al.,^27^ and the modifications applied in this study are given in the Supporting Information. In brief, the model is based on a spherical shell diffusion system. However, the droplets in the present work are large (millimeter-sized) and not completely spherical, but to some degree spread out in the wells. Hence, consideration of their flattened shape is required. This is done by describing the flattened droplet with average thickness *h* as a hemispherical shell with equal thickness concentrically covering an inert core (see **Figure S4**). To inform the model, IAV inactivation was first parameterized in terms of the NaCl molality and sucrose content. Generally, while the conditions in the droplets change as a function of time, one cannot expect first order inactivation kinetics. Conversely, we assumed that droplets exhibiting first-order inactivation are in equilibrium with the surrounding air. This can occur, for instance, after efflorescence or when the viscosity of the droplets becomes high and the diffusivity within them is low, such that their salt molality no longer changes.

The dependence of inactivation on NaCl molality was determined based on data obtained in both droplet and bulk experiments. Specifically, the molality of any residual fluid in effloresced NaCl droplets following exposure to 30% RH corresponds to saturation molality, i.e. 6.1 mol NaCl/kg H_2_O = 6.1 molal,^48^ whereas the molality in deliquesced droplets at 65% RH corresponds to a supersaturated state of ca. 8 mol/kg. In agreement with this molality difference, the IAV inactivation rate coefficient *k* in the residual fluid at 30% RH (after efflorescence) was lower than that measured at 65% (**Figure 2** and **Table S2**). To establish a relationship between NaCl molality and *k* over a wide molality range, we furthermore measured inactivation in bulk solutions of 0.14, 2.8 and 6.1-mol/kg NaCl (**Figure S5)**. The resulting time courses were well fit by a first-order model, despite a short initial period with faster decay. Finally, the rate coefficient for saturated salt solutions (**Table S2**) was confirmed in a droplet experiment conducted in NaCl at 74% RH (**Figure S6**), where the NaCl molality in the equilibrated droplet corresponds to 6.1 mol/kg. Based on these data, the following exponential relationship between NaCl molality and inactivation rate *k* (in s^-1^) and corresponding e-folding time *τ* could be established (**Figure 4B**):

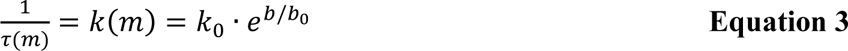

**Figure 4:**
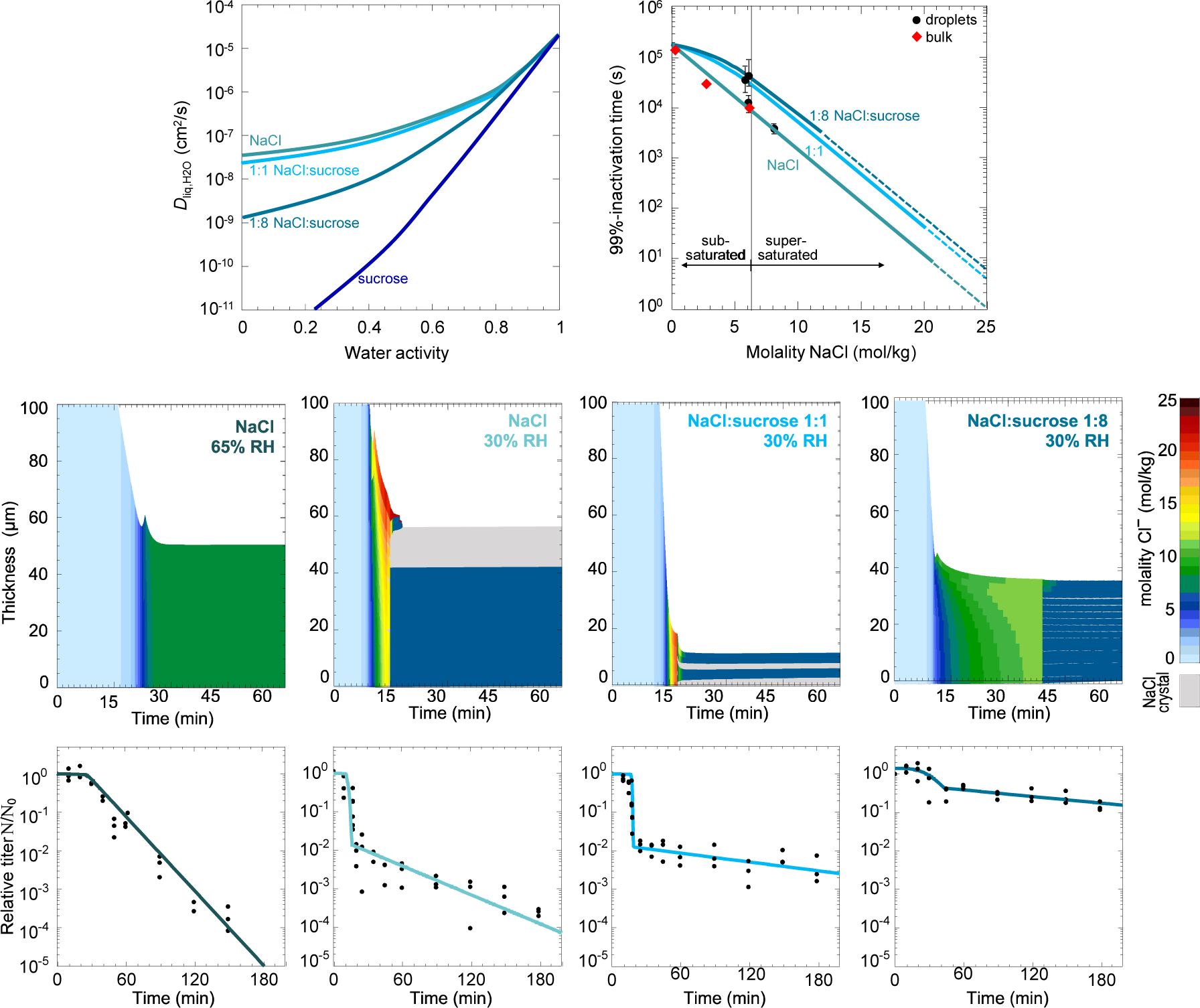
IAV inactivation in deposited droplets modeled with ResAM. (A) Liquid phase diffusion coefficients of H_2_O molecules in aqueous NaCl and sucrose solutions and 1:1 and 1:8 dry mass ratio mixtures thereof (see **Equations S1 to S3**). (B) Time required for 99% titer reduction of IAV versus NaCl molality in aqueous NaCl/sucrose droplets derived by fitting ResAM to the experimental data in panels (G-J) and in bulk NaCl solutions. The inactivation times in droplets are based on the first order inactivation rates after droplet compositions stopped changing by equilibration with air, that is ≥ 40 min after deposition. The green line represents the model prediction of 99%-inactivation time according to **Equation 3** for NaCl only. The light blue and dark blue lines are for aqueous NaCl:sucrose mixtures with a dry mass ratio of 1:1 and 1:8, respectively, according to **Equation 4**. (C-F) Evolution of NaCl molality in 1-μl drying droplets with different compositions, modeled with ResAM. At 30% RH, all the droplets reach a final molality of 6.1 mol/kg, corresponding to saturation conditions after efflorescence independent on sucrose content. Light gray regions show the effloresced NaCl crystals, which are modeled as dendrites with average distances of 5 um, as observed by Luo et al.^27^ in synthetic lung fluid. In (D) a crust of pure salt hinders the further diffusion of H_2_O, whereas in (E,F) the (assumed) salt layers interspersed with sucrose reduce the aqueous NaCl concentration but remain penetrable for H_2_O molecules. (G-J) Measured IAV inactivation in 1-μl droplets (black dots) in NaCl at 65% RH (G), NaCl at 30% RH (H), NaCl:sucrose 1:1 at 30% RH (I) and NaCl:sucrose 1:8 at 30% RH (J) and simulated inactivation curve (solid lines) determined using ResAM.

where *b* is the molality of NaCl (in mol/(kg H_2_O)), *b*_0_ is a fitting parameter corresponding to 2.095 mol/(kg H_2_O); and *k*_0_ is the fitted inactivation rate coefficient in pure water, corresponding to 2.5·10^-5^ s^-1^.

For sucrose-containing droplets, a similar exponential dependence of *k* on NaCl molality was assumed, though the absolute value of *k* at saturation molality was lower due to the protective effect of sucrose exerted in equilibrated droplets (**Figure 3B**). To account for the effect of sucrose, the relationship shown in **Equation 3** was expanded as follows:

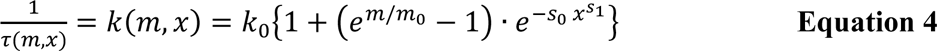

where *x* is the mole fraction of sucrose defined as *x* = *m*_sucrose_/(*m* + *m*_sucrose_), and *s*_0_ = 1.8696 and *s*_1_ = 0.19451 are fitting parameters. For *x* = 0, i.e. in the absence of sucrose, **Equation 4** reduces to **Equation 3**. The inactivation rate in pure water is *k*_0_, while the impact of salt is given by the term (*e^m/m_o_^* – 1) and protection effect by sucrose by the term *e*^−*s_o_ x*^*s*_1_^^. The fitting parameters in **Equations 3** and **4**, *k*_0_, *b*_0_, *s*_0_ and *s*_1_, were determined by optimizing the output of ResAM to match the measured titers in the NaCl-sucrose solutions, see **Figure 4G-J**. This fitting procedure must also properly account for the diffusivities of H_2_O molecules in these solutions (see **Figure 4A**). While appearing arbitrary at first glance, these fitting parameters are actually quite well constrained for water activities *a_w_* > 0.74.^49^ Mapping the kink in the curves just before efflorescence (**Figure 4G-J**) is a challenge that effectively limits the fitting options (calculations not shown).

Next, ResAM was used to model the water activity and the NaCl molality in the different NaCl droplets. Hereby some assumptions were made: because the inactivation rate coefficient observed in the tail of the pure salt droplet at 30% RH corresponds to that of saturated NaCl solutions (as measured in bulk and in droplets at 74%; see **Figure 4B** and Supporting Information), we conclude that NaCl crystals grow around the virus, forming pockets that limit water evaporation and in which the aqueous salt is saturated. Furthermore, while the 1:8 NaCl:sucrose matrix does not show efflorescence visible by eye, the observed reduction in the inactivation rate after 45 min is indicative of efflorescence. We therefore assume that microscopic dendritic crystal growth occurred in this matrix. This assumption is needed to ensure that the model is consistent with the experimental data. Finally, the model assumes a homogeneous distribution of the virus in the droplet. This is in contrast with studies suggesting that the virus may preferentially localize on the border of the dried droplet when proteins are present in the matrix.^24,39^ The exact location of IAV viruses in a drying NaCl/sucrose droplet remains to be assessed.

Under these assumptions, ResAM revealed the following: (i) at 65% RH, (**Figure 4C**) the NaCl molality increases up to 8 mol/kg, until equilibrium with air. No efflorescence occurs at this RH, such that the droplet remains supersaturated in NaCl; (ii) at 30% RH (**Figure 4D**), NaCl molality first increases due to water evaporation, and reaches supersaturation (up to 20 mol/kg at the particle surface) before the salt effloresces. After efflorescence, the concentration drops down to saturation molality (i.e., 6.1 mol/kg) and inactivation continues with the same rate as in a saturated bulk solution, likely in pockets of solution encapsulated by crystalline salt; and (iii) in sucrose-containing droplets at 30% RH, the NaCl molality prior to efflorescence is dependent on the sucrose content: the more sucrose, the lower the diffusivity of water and hence the greater the retention of water inside the droplet. (**Figure 4D-F**). After efflorescence, any remaining liquid is in equilibrium with crystalline salt, and hence the aqueous NaCl molality in all matrices corresponds to saturation, i.e. 6.1 mol/kg. The water activities within the drying droplets are shown in **Figure S7.** The diffusion coefficient of water in the different NaCl:sucrose matrices are shown in **Figure 4A**.

Finally, by combining the simulated NaCl molality and the molality- and sucrose-dependent inactivation rate coefficient, IAV inactivation was calculated in each shell and at each time point, and the residual virus concentration was integrated over the entire droplet. The resulting inactivation curves correspond well to the measured data in all droplets investigated (**Figure 4G-J**). The model was able to capture the inverted sigmoidal curve of the inactivation in effloresced droplets, the first-order inactivation in deliquesced droplets, and was able to reproduce the protective effect of sucrose.

### Effect of NaCl on infection cycle and virion integrity

To gain insight into the mechanism of NaCl-induced inactivation, we first conducted fluorescence imaging of viral replication within cells (see Supporting Methods). This is done by fluorescently labeling the influenza nucleoprotein (NP), which is expressed within the host cell’s nucleus during the early stages of IAV infection cycle, and is then transported in the cytoplasm at a later stage.^50^ Therefore, the presence of NP fluorescence within the nucleus or the cytoplasm of the cells indicates active viral replication, while no signal indicates the virus’s incapacity to synthesize NP – i.e., the virus is inactivated. For this purpose, we used sample aliquots from the inactivation experiment in 1-μl NaCl droplets exposed to 30% or 65% RH (**Figure 2B**). At both RH levels, the IAV titer was reduced by ∼3-log_10_ after 120 min. Samples taken at times 0 and 120 min were used to infect A549 cells for 6 h, prior to fluorescent staining of the IAV nucleoprotein (NP). A549 cell nuclei were stained with DAPI. Fluorescence microscopy showed that in cells infected with viruses in freshly deposited droplets (exposure time 0 min), NP production was present in approximately half of the cells (**Figure 5A**), indicating that viral replication occurred. However, viruses exposed to 30% or 65% RH for 120 min did not show any NP signal 6 h post infection, indicating that the infection cycle was interrupted at an early stage (that is, before the onset of NP protein synthesis).

**Figure 5:**
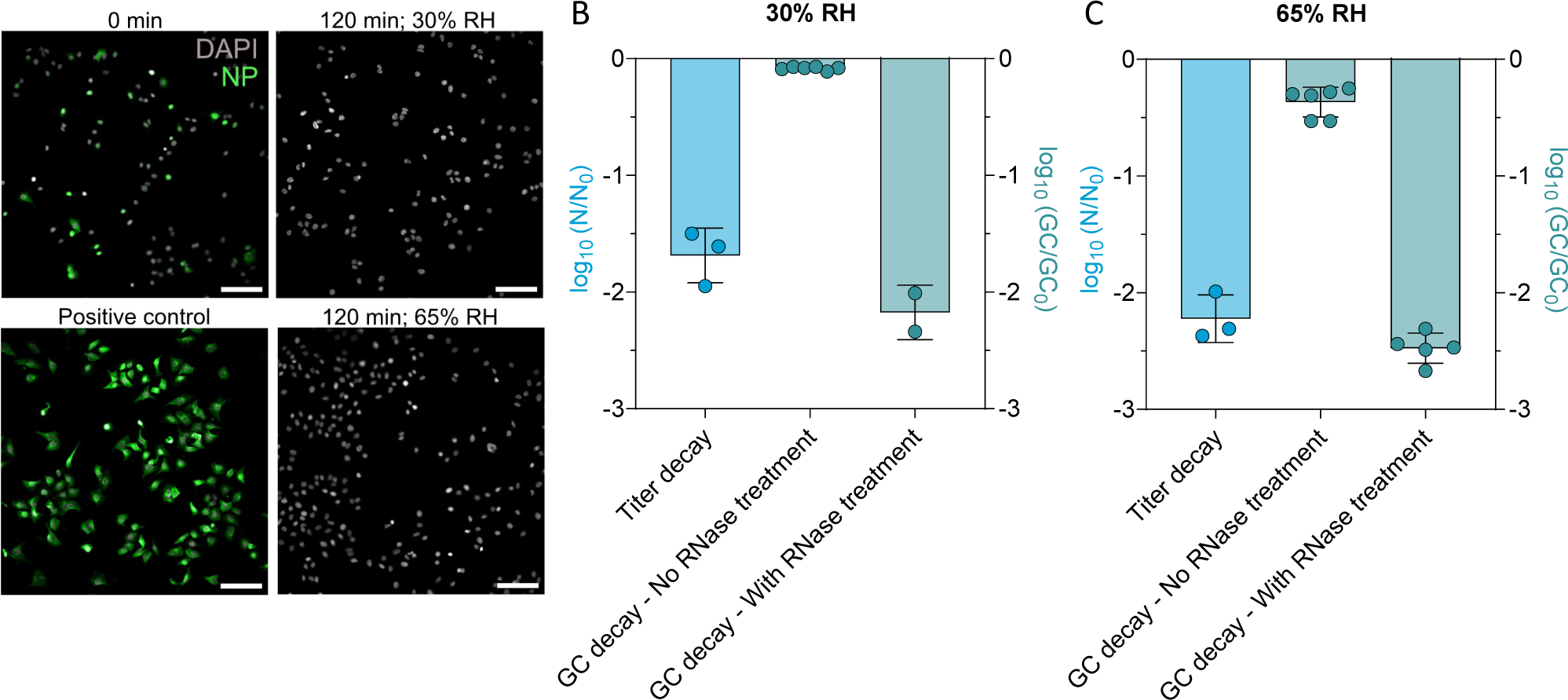
Effect of high salinity on IAV infectivity. (A) Immunofluorescence images of A549 cells infected with IAV samples exposed to 30% or 65% RH in 1-μl NaCl droplets for 120 min. A549 cells were infected at a multiplicity of infection (MOI) of 1, and IAV protein production was visualized by immunofluorescent staining of nucleoprotein at 6 h post-infection. Scale bar corresponds to 100 µm. Images are representative of duplicate experiments. The positive control was infected with a fresh stock of IAV. (B,C) Infectivity decay of IAV samples exposed to 30% (B) or 65% RH (C) in 1-μl NaCl droplets for 120 min, and genome quantification by RT-qPCR of these samples incubated with and without RNase for 30 min prior to RNA extraction. Data are presented as the log_10_ of the ratio between viral concentration after 120-min exposure in droplet and the initial concentration. In case of infectivity measurements, the virus concentration is represented by the titer (in PFU/ml), and in case of RT-qPCR quantification, the virus concentration is represented by the genomic copies (in GC/ml).

Next, we investigated the integrity of the virions after droplet drying (see Supporting Methods). To this end, 1-μl NaCl droplets were again exposed to 30 and 65% RH, resulting in 1.5-2.5 log_10_ inactivation after 120 min (**Figure 5B-C**). Samples taken at times 0 and 120 min were then incubated for 30 min at 37°C in a digestion mixture (see composition in *Methods*) containing RNase, and the residual RNA copy number was subsequently quantified by RT-qPCR. Hereby the assumption is that if the virion integrity is compromised by exposure to NaCl, the RNase can access and degrade the viral RNA, whereas in intact virions, the RNA is protected.^51^ As a control, half the volume of each sample was exposed to the digestion mixture without RNase. In the absence of RNAse, only a minor loss of genome copies was observed over the course of 120 min. In contrast, in the presence of RNAse, approximately 2-log_10_ decrease in genomic copies was observed after 120 min at both humidities (**Figure 5B-C**, third bar). This decay corresponds well to the observed infectivity decay (**Figure 5B-C**, first bar), suggesting that the disruption of virion integrity is the main inactivation mechanism at high NaCl concentrations.

## DISCUSSION

Salinity, especially the presence of NaCl but also of other salts such as LiCl and KCl, has been proposed as a promoter for the inactivation of viruses,^52–54^ yet a comprehensive understanding of the underlying mechanisms and kinetics has remained elusive. In this study, we combined experimental data on IAV inactivation at different salinities with a biophysical model to parameterize the effect of NaCl on virus stability. Our data suggest that the main inactivation driver of IAV in efflorescing saline droplets is exposure to high (supersaturated) NaCl molality rather than the formation of salt crystals during efflorescence. This suggestion was confirmed by model simulations showing that the NaCl molality, which increases dramatically prior to efflorescence and then is reduced to saturation post-efflorescence, is sufficient to explain the observed inactivation.

The ResAM model was able to capture the entire inactivation curve observed in both effloresced and deliquesced droplets, and in both aqueous NaCl and NaCl/sucrose matrices. A recent study proposed the inactivation of bacteriophage MS2 to be correlated with the cumulative dose, defined as the solute concentration multiplied by the exposure time.^18^ A dose-dependent inactivation could not be confirmed by ResAM for IAV. Instead, we here demonstrate an exponential dependence of IAV inactivation on salt molality, suggesting that an increase in molality leads to a disproportionate increase in inactivation. In particular at salt molalities > 20 mol/kg, inactivation times become very fast, with 99% inactivation achieved within seconds. However, even at a RH as low as 30%, it takes several minutes to reach such molalities in 1-μl NaCl droplets, and in droplets containing large amounts of organics, such molalities may not be reached at all. Depending on the RH and the organic content, the virus may thus remain stable for minutes to hours. Therefore, droplet transmission may play an important role in the spread of IAV, and a combination of prevention strategies addressing both aerosol (e.g. ventilation) and droplet transmission (e.g. surface disinfection), may be needed to prevent the spread of influenza.

While our results were obtained in millimeter-sized deposited droplets, our findings are also relevant to smaller, airborne particles with diameters up to few micrometers. Below the deliquescence RH (74%), such particles can rapidly reach supersaturated salt concentrations, similar to those obtained in our experimental system (but experimentally inaccessible in bulk solutions). As shown in our experiments with sucrose, supersaturation is also achieved in matrices with high amounts of organics. We therefore expect that the observed dependence of IAV stability on NaCl molality also applies to smaller particles, including organic-rich expiratory aerosol particles. Depending on the composition of the respiratory matrix and the surrounding air, however, the inactivating effect of NaCl in such small particles may be outcompeted by other inactivating factors, including aerosol pH. Specifically, virus inactivation in small particles was previously suggested to be driven by acidic^27^ or basic^29^ pH, though the contribution of pH-mediated inactivation mechanism remains to be confirmed in real world systems. In contrast, acidification in millimeter-sized droplets is too slow to compete with salt-mediated inactivation,^27^ and we found IAV to be stable over the course of 1 h at an alkaline pH of 11 (**Figure S8**). We can therefore exclude pH as a determinant of virus inactivation in droplets, highlighting the importance of salinity in modulating virus stability.

The inactivation of IAV in drying droplets may be explained by a combined effect of two processes: the removal of water molecules due to evaporation, and the subsequent increase of salt molality. Our data suggest that the virion integrity is damaged in drying NaCl droplets (**Figure 5**). Virion integrity loss was also observed to be a minor mechanism in the inactivation of IAV by acidic pH, as encountered in small aerosol particles.^51^ For salinity-mediated inactivation in drying droplets, however, damage to the virus structure seems to be the main inactivation mechanism. Therefore, we here summarize how these two processes may affect the lipid envelope and the viral proteins.

Upon drying, the osmotic pressure induced by the solutes increases and causes shrinkage of IAV by forcing water molecules to exit the virus.^55^ The shrinkage rate constant of influenza virus was previously observed to increase with ionic strength, until reaching a threshold where the lipids and the matrix proteins (M1) are no longer able to adapt to the compressive stress, resulting in morphological change and loss of membrane integrity.^55^ This concurs with our finding that damage to the viral integrity is the main inactivation mechanism. The authors furthermore hypothesize that the morphological change in the membrane leads to irreversible conformational changes, resulting in the loss of hemagglutinin (HA) activity. We did not investigate the structural integrity and functionality of HA in this work, though it is possible that this mechanism occurs concomitantly to the loss of virion integrity in drying saline droplets.

In addition to osmotic stress, water evaporation may induce leakage of the viral lipid envelope. It was shown that when the water molecules surrounding the phosphate heads of membrane phospholipids are removed, the packing of the head groups densifies. The subsequent increase of Van der Waals’ interactions among hydrocarbon chains trigger a phase transition in the lipid bilayer.^56,57^ Specifically, under physiological conditions, the phase transition temperature, (T_m_) is usually below room temperature, and the hydrated lipids are in a disordered, flexible liquid crystalline phase. However, the increase of Van der Waals’ interactions due to water removal increases T_m_ to values above room temperature and forces the dried lipids to transition into an ordered, solid-like gel phase. Upon rehydration, the lipids then undergo a reverse phase transition when the head group packing decreases due to the formation of H-bonds with water. These phase transitions induce leakage of the lipid bilayer, damaging the membrane. We hypothesize that a similar mechanism occurs in the viral envelope of influenza virus in a drying droplet or aerosol particle, with rehydration occurring when the particle is inhaled by a host. Moreover, both Na^+^ and Cl^-^ ions were found to bind with the lipid head groups, increasing the ordering of the lipids and subsequently increasing T_m_.^58–61^ We propose that the combination of water removal and NaCl attachment to the lipid bilayer promote phase transition and cause damage to the viral envelope, which may interfere with fusion during viral infection. Such a mechanism agrees with our data that demonstrate that in drying droplets, IAV inactivation is driven by high NaCl molality and is caused by damaging the virion integrity. It is also consistent with findings by Lin et al.,^62^ who observed a higher salt susceptibility of the enveloped bacteriophage Φ6 than the non-enveloped bacteriophage MS2, supporting the hypothesis of viral envelope disruption by high salinity. Similarly, Kukkaro and Bamford^63^ saw a larger sensitivity to NaCl concentration changes of enveloped viruses compared to non-enveloped ones. Nevertheless, to conclusively confirm this mechanism, future work should study phase transition mechanism in the lipid bilayers of enveloped viruses in drying droplets.

We observed that the addition of sucrose is protective for IAV in drying droplets (**Figure 3B**). The protective mechanism by sucrose is two-fold. First, as shown in **Figure 4**, the presence of sucrose in the matrix decreases the molality of NaCl during the drying phase, thereby lowering viral inactivation. This effect is particularly evident at an NaCl:sucrose ratio of 1:8, and constitutes an indirect protective effect from sucrose. Second, our results show that sucrose also has a direct protective mechanism: in droplets at equilibrium, the NaCl molality is independent of the sucrose concentration (**Figure 4B**), and yet lower inactivation rate coefficients are observed when sucrose is present (**Table S2**). The inactivation rate coefficients follow the trend 1:0 > 1:1 > 1:8 (**Figure 4H-J**), showing that an increasing sucrose concentration increases the protection. Here, the protection cannot be attributed to a change in NaCl molality, as the latter is at saturation in all conditions. Therefore, a different, direct mechanism must protect the virus.

Potential direct protection mechanisms by sucrose acting on both lipids and proteins are described in the literature. The addition of disaccharides, such as sucrose and trehalose, was previously shown to help stabilize the membrane phospholipids during lyophilization.^56,57,64^ The sugar molecules interact with the phosphates, inserting in between the head groups and replacing water. The packing density needs to decrease to give room to the sugars, and the Van der Waals’ interactions are reduced. Consequently, T_m_ is decreased, and remains below room temperature. Therefore, despite water removal, the lipids do not undergo phase transition, and no membrane leakage occurs. The direct protection of IAV exerted by sucrose in the droplet may thus be partly due to the protection of the lipid bilayer from water loss, preventing the damage to the lipid envelope during drying. In addition, the sucrose may also protect the lipid bilayer from the effects of salt, if sucrose replaces the Na^+^ and Cl^-^ ions between the lipid head groups and thereby counteracts the increase in T_m_ caused by the salt. This mechanism remains to be assessed, but it correlates with the decrease in inactivation rate as a function of sucrose content in matrices with similar NaCl molalities (**Figure 4H-J**).

Additionally, protection may arise from the stabilization of surface proteins by sucrose.^65–68^ Upon drying, the water molecules stabilizing the structure of proteins are removed, unfolding the proteins and inducing potentially irreversible conformational changes.^66,69^ However, when osmolytes (here, sucrose) are present, they are sterically excluded from the first hydration shell of the proteins. This preferential exclusion of the sucrose leads to a preferential hydration of the proteins, increasing the free energy of the system. The unfolded state of the proteins becomes thermodynamically unfavorable, forcing the proteins to remain in their native state. Our data do not indicate whether protein unfolding occurred in drying droplets, and therefore we cannot assess whether the sucrose protection is in fact caused by protein stabilization. Indeed, the surface proteins of a virus have lower mobility than proteins in solution, and it is possible that this stabilization mechanism is less pronounced in viruses. Further studies on IAV are required to confirm this protection mechanism by sucrose and other organics present in respiratory fluids.

This study has some limitations. First, the ResAM model used herein relies on some assumptions that remain to be validated. Specifically, the encapsulation of residual water and viruses by NaCl crystals in efflorescing droplets, as well as the formation of micro-crystals in the 1:8 NaCl:sucrose solution, are hypothetical physical processes that need to be assessed. However, previous studies have shown that the viscosity of an organic phase can kinetically limit the coalescence of the inorganic salt crystal,^70^ and non-coalesced salt crystals were observed in porcine respiratory fluid,^71^ which supports the hypothesis of micro-crystal formation when the sucrose content is large. Moreover, we accounted for the flattened shape of the droplet despite using a spherical model by limiting the diffusion depth to the thickness of the spread droplet. This also represents a minor uncertainty of ResAM, as detailed in the SI. The likely largest uncertainty in ResAM, however, results from the extrapolations of the inactivation times (**Figure 4B**) and the diffusivities (**Figure 4A**) to highly supersaturated regimes without direct measurements. For both these parameters, we conducted sensitivity analyses (see SI and **Figures S9** and **S10**), which reveal that in particular, improved experimental data for diffusivities in highly concentrated salt solutions would be beneficial for further constraining our model assumptions.

Finally, the simple NaCl/sucrose/water matrices that were used in this study are useful to parameterize ResAM, but are far less complex than real respiratory fluids and saliva. In real matrices, other processes may occur during droplet drying, such as the formation of a glassy shell by proteins, limiting evaporation.^32^ In addition, the morphology of the dried droplet, the importance of which has been recently presented by Rockey et al.,^39^ is dependent on the matrix composition, such as the Na:K ratio^71^ and the initial NaCl content^34,35^ that can affect crystal formation. It is therefore essential to combine studies of both simple and more complex matrices, to disentangle the effects of all parameters and assess their importance for IAV transmission.

In summary, we demonstrate that IAV inactivation in deposited droplets is driven by the high NaCl molality due to water evaporation. Furthermore, we propose a model to simulate IAV inactivation in droplets as a function of RH, NaCl and organic concentration. Finally, we determined that organics (sucrose) protect IAV by two mechanisms: first by decreasing the salt molality in the drying droplet and second, by a direct mechanism that may prevent the leakage of the lipid bilayer and the unfolding of surface proteins. This work contributes to the understanding and quantification of virus inactivation in drying droplets. It expands the fundamental knowledge of airborne virus transmission, which is the basis for the development of appropriate mitigation strategies.

## ASSOCIATED CONTENT

### Supporting information

The following file is available free of charge and contains:

Method descriptions for virus propagation, purification and quantification, nucleoprotein staining by immunofluorescence and RNase assay to test virion integrity; further description of ResAM adaptation to droplets; discussion of ResAM limitations; droplet pictures; additional IAV inactivation kinetic measurements in bulk and in droplets; check-list for RT-qPCR following MIQE guidelines; values of inactivation rate coefficients and corresponding 99%-inactivation times (PDF)

## AUTHOR INFORMATION

### Corresponding Author

Tamar Kohn –Laboratory of Environmental Virology, School of Architecture, Civil & Environmental Engineering, École Polytechnique Fédérale de Lausanne (EPFL), CH-1015 Lausanne, Switzerland –

### Author Contributions

Conceptualization: AS, TK, TP, UKK

Methodology: AS, BL, IG, LKK, SCD, TK, TP, UKK

Software: BL

Formal analysis: AS, BL, TP

Investigation: AS, BL, IG, LC, SCD

Resources: UKK, SSt, TK

Writing–original draft: AS, BL, TK, TP

Writing–review & editing: AS, BL, CT, GM, IG, KV, LKK, MP, NB, SCD, SSt, TP, TK, UKK

Visualization: AS, BL, IG, LKK, TK, TP

Supervision: SSt, TK, TP, UKK

Project administration: TK

Funding acquisition: AN, SSt, TK, TP, UKK, WH

### Notes

Authors declare that they have no competing interests.

### Data availability

Experimental data are available in the public repository Zenodo under the following link: https://zenodo.org/records/10418730. ResAM code will be made available upon manuscript acceptance.

## ACKNOWLEDGEMENTS

This work was funded by the Swiss National Science Foundation (grant 189939). The authors thank Sophia Kiselova for assistance with the illustrations.

