## Supplementary Material for "Salt supersaturation as accelerator of influenza A virus inactivation in 1-μl droplets"

**This PDF file includes:**

22 pages
Supporting Experimental Methods
Biophysical modelling: further description of ResAM adaptation to droplets
Model limitations
Figure S1 to S10
Tables S1 and S2
References

### Supporting Experimental Methods

#### Virus propagation and purification

Experiments were conducted with influenza A virus (IAV) strain A/WSN/33 (H1N1 subtype). Madin-Darby Canine Kidney (MDCK) cells (ThermoFisher) maintained in Dulbecco's modified Eagle medium (DMEM, Gibco) supplemented with 10% Fetal Bovine Serum (FBS; Gibco), and 1% Penicillin-Streptomycin 10,000 U/ml (P/S; Gibco) were used for propagation. Confluent MDCK monolayers were washed and inoculated with IAV at a multiplicity of infection (MOI) of 0.001 for 72 h in OptiMEM (Gibco) supplemented with 1% P/S and 1 µg/ml TPCK trypsin (Sigma, T1426). The culture supernatants from the infected cells were cleared by centrifugation (2,500 ×g, 10 min). Subsequently, the viruses were concentrated by ultracentrifugation by pelleting through a 30% sucrose cushion at 112,400 ×g in a SW31Ti rotor (Beckman) (90 min, 4°C). The pellets were resuspended in phosphate-buffered saline (PBS, ThermoFisher, 18912014) overnight at 4°C. Concentrated IAV stock solutions were quantified by plaque assay (described below) resulting in a titer of ~$10^{11}$ plaque forming units (PFU)/ml. The stock solution was aliquoted and frozen at -80°C until use.

#### Quantification of IAV infectivity

Plaque assay was performed by infection of MDCK cells, as described in Luo et al.[1] Briefly, a 12-well plate containing MDCK monolayers was washed with PBS and infected with IAV samples. Prior to infection, the samples were serially diluted in PBSi (PBS for infection; PBS supplemented with 1% P/S, 0.02 mM $Mg^{2+}$, 0.01 mM $Ca^{2+}$, and 0.3% bovine serum albumin (BSA, Sigma-Aldrich A1595), with a final pH of ~7.3). A negative control was always performed on an additional plate, using PBSi as a blank sample. Infected monolayers were incubated for 1 h at 37°C with 5% of $CO_2$, and manually agitated every 10 min. The inoculum containing non-attached viruses was removed and the cells were covered by an agar overlay (MEM supplemented with 0.5 µg/ml of TPCK-trypsin and 2% of oxoid agar (Thermo Fischer LP0028-500G)). After 72 h of incubation (37°C, 5% $CO_2$), cells were fixed (PBS + 10% formaldehyde (Sigma, 47608-1L-F)) and stained (0.2% crystal violet solution (Sigma, HT901-8FOZ) in water + 10% methanol (Fisher Chemical, M-4000-15)) to enumerate and determine the virus titer in PFU/ml.

#### Quantification of IAV genome copies

RNA extractions were performed using the QIAamp Viral RNA Mini extraction kit (Qiagen, 52906) according to manufacturer's instructions. Sample aliquots of 140 µl were extracted and nucleic acids were eluted in 80 µl of elution buffer, and stored at -20°C until analysis. A negative extraction control was always performed to ensure the clean state of the kit columns and reagent. Amplification and detection were performed using the One Step PrimeScript™ RT-PCR Kit (RR064A, Takara Bio) with the following primers targeting a 244-base amplicon of the IAV M segment:

Forward primer 5'-ATGAGYCTTYTAACCGAGGTCGAAACG-3'
Reverse primer 5'-TGGACAAANCGTCTACGCTGCAG-3'

The reaction mixture used for the one-step Reverse Transcription quantitative Polymerase Chain Reaction (RT-qPCR) assay was composed of 7.5 µl of 2x One-Step SYBR RT-PCR Buffer, 0.3 µl of Takara Ex Taq HS (5 U/µl stock), 0.3 µl PrimeScript RT enzyme mix, 0.3 µl forward and reverse primers (10 µM stocks), and 3.3 µl RNase-free water, to which 3 µl extracted RNA were added. RT-qPCR was performed using a Mic Real-Time PCR System from Bio Molecular Systems. The run profile was the following: 2 min at 50°C and 10 min at 95°C for reverse transcription and denaturation, followed by 40 cycles of 15 sec at 95°C and 60 sec and 60°C for annealing and extension, and a final dissociation step from 55°C to 95°C at 0.3°C/s. A Gblock gene fragment, as described in [2], was used to create a standard curve for quantification over a range of $10^2$ to $10^7$ GC/µl . A no-template control of milli-Q water was included in every run and was always negative. The absence of inhibition was regularly checked using serial dilutions of the samples. The micPCR software (version 2.12.16) was used to acquire the qPCR data. The Generic qPCR Limit of Detection (LOD) / Limit of Quantification (LOQ) calculator [3] was used to analyze pooled standard curves. The average slope of the standard curve was -3.47 and the average intercept was 34.22, with a $R^2$ of 0.999 and a PCR efficiency of 93.2%. The limit of quantification was determined at 100 copies/reaction. The LOQ is defined as the lowest standard concentration with a coefficient of variation smaller 35%. All RT-qPCR procedures followed MIQE guidelines (see **Table S1**).

**Nucleoprotein (NP) staining by immunofluorescence**

Droplets were recovered from exposure to 30% and 65% RH as described in the methods section. For each RH, one droplet was recovered immediately, and two droplets were recovered after 120 min. Each experiment was performed in duplicate. Samples were frozen at -20°C until analysis. A549 cells were infected with the samples at an MOI of 1 and incubated at 37°C for 1 h. The cells were then rinsed with PBS and incubated for 2 or 5 h in DMEM supplemented with P/S, 0.2% BSA, 20 mM HEPES (Sigma-Aldrich, H7523), and 0.1% FBS. After incubation, 3.7% formaldehyde (ThermoFisher Scientific, 26908) in PBS was used to fix the infected cells for 15 min, followed by blocking and permeabilization for 1 h at room temperature in PBS containing 50 mM ammonium chloride (Sigma-Aldrich, 254134), 0.1% saponin (Sigma-Aldrich, 47036), and 2% BSA (Sigma-Aldrich, A7906). Subsequently, a mouse monoclonal anti-NP antibody (hybridoma supernatant, ATCC, HB-65) was applied for 1 h at room temperature. After washing the cells with PBS, the secondary antibody, anti-mouse IgG Alexa488 (ThermoFisher Scientific, A-11029), was added, as well as DAPI (Sigma-Aldrich, 10236276001) to stain the nuclei. After 1 h of incubation at room temperature, the cells were washed and cover slips were mounted with ProLong Gold Antifade Mountant (Thermo Fisher Scientific, P36930). The DMi8 microscope (Leica) in combination with the THUNDER Instant Computational Clearing algorithm (Leica) were used to image the stained samples.

The detection range of this assay depends on the number of cells in the imaging window; this number is ~100, therefore the maximum inactivation that can be detected is ~2-$\log_{10}$. In comparison, the infectivity quantification by plaque assay in our experimental system extends over a range of ~5-$\log_{10}$. Inactivation exceeding 2-$\log_{10}$ can thus be measured by plaque assay but give negative results by NP staining.

**RNase assay to test virion integrity**

Droplets were recovered from exposure to 30% and 65% RH as described above. For each RH, three droplets were recovered immediately, and three droplets were recovered after 120 min. Infectivity was quantified for each sample. From each sample, 1.5 µl aliquots were diluted in RNase digestion mixture to a total volume of 150 µl, in duplicate. The mixture was composed of TE buffer containing 10 mM Tris and 1 mM EDTA (Invitrogen, AM9858) supplemented with 50 mM NaCl. The pH was adjusted to 7.5 with HCl. One replicate of each sample was supplemented with RNase A/T1 (ThermoFisher Scientific, EN0551; final concentration was 40 µg/ml RNase A and 100 U/ml T1. The other replicate was supplemented with an equal volume of PBS and served as the RNAse-free control. All samples were vortexed and incubated for 30 min at 37°C. After incubation, 1 µl of SUPERase-In RNase Inhibitor (Invitrogen, AM2694, acts against RNase A, B, C, 1, T1) was added to each sample, and a second incubation (room temperature, 20 min) was performed before freezing the samples at -20°C for at least one night prior to RNA extraction.

#### Biophysical modelling: further description of ResAM adaptation to droplets

The data were modeled using the Respiratory Aerosol Model ResAM, fully described by Luo et al.[1] Briefly, a spherical shell diffusion model is used to determine viral inactivation in droplets, considering their initial composition and size, as well as the relative humidity (RH), temperature and ventilation in the experimental chamber, assuming typical indoor air without trace gases. Water activity, sodium chloride and sucrose concentrations, as well as the components of PBS within the droplet are simulated as a function of time, taking into account the full composition with $H_2O$, $H^+$, $OH^-$, $Na^+$, $Cl^-$, $K^+$, $HPO_4^{2-}$, $H_2PO_4^-$, $H_3PO_4$, $CO_3^{2-}$, $HCO_3^-$, $CO_{2,aq}$, and sucrose (where the phosphorous species are new in this model version with dissociation constants[4] and the activity coefficients of sulfate ions and $K^+$ were treated like $X^+$ in Luo et al.[1]). A description of the spherical shell modeling, as applied to airborne exhalation droplets, is given by Luo et al.[1] Here, however, the droplets are not suspended in air and are thus not spherical, but are placed on the bottom surface of the wells. These have been treated such that under aqueous assay conditions, a hydrate layer forms that prevents dissolved biomolecules from binding to the microplate surface (Greiner Bio-One, 655901), with the disadvantage that droplets tend to partially wet the surface with a contact angle $\alpha < 90°$. Assuming the droplet shape to be a spherical cap and using the video images and the initial volume of 1 µl, we estimate their initial contact angle as $\alpha \approx 40°$ (see top row in **Figure S1**). Furthermore, the treatment of solid crystal formation has been refined compared to Luo et al.[1] by not assuming a concentric salt core but permeable shells with a spacing of 5 µm, corresponding to the characteristic distance of the salt dendrites shown in Figure S6 of Luo et al.[1]

Accurate modeling of the aspherical droplet geometry on the bottom of the well and the diffusion of $H_2O$ molecules in the cylindrical shape of the well would be challenging and not worth the effort. Rather, it is important to correctly capture the physicochemical changes and the viral inactivation in the droplets, which are determined by the transport velocity of the $H_2O$ molecules as they diffuse through the millimeter-thick liquid matrix towards the droplet surface and from there via the gas phase in the well into the free gas space of the environmental chamber. Therefore, in order to describe the composition change and IAV inactivation in the droplets, it is crucial to correctly describe these diffusion processes. To this end we use ResAM with concentric shells, having converted the ellipsoidal shape of the large, millimeter-sized droplets into the geometry of a hemispherical shell residing on an inert solid core that sets the correct average thickness of the droplet, see **Figure S4**. Except for the edge regions of the droplet, where the approximation of radial symmetry fails, this is a very good description of the diffusion processes for most of the droplet volume.

The diffusion coefficients of $H_2O$ molecules in binary aqueous sucrose solution, $D_{\mathrm{H_2O,sucrose}}$, have been taken from Zobrist et al.[5] For binary aqueous NaCl solutions, the concentration dependence of $H_2O$ diffusivity is obtained from viscosity data obtained by Kestin and Shankland[6] for solutions of $b$ < 6.1 mol kg$^{-1}$ (i.e., water activity > 0.74) using the Stokes-Einstein relation:

$$ln\left(\frac{D_{\mathrm{H_2O,NaCl}}(mb)}{D_{\mathrm{H_2O}}(0)}\right) = ln\left(\frac{\eta(0)}{\eta(mb)}\right)$$
$$= -(0.21319213 + 0.13651589 \times 10^{-2}\, T - 0.12191756 \times 10^{-5}\, T^2) \times b$$
$$-(0.069161945 - 0.27292263 \times 10^{-3}\, T + 0.20852448 \times 10^{-6}\, T^2) \times b^2$$

**Equation S1**

Here, $b$ is the molality and $\eta$ is the viscosity of the solution and $T$ the temperature in K.

For water activity < 0.74, which is difficult to access in bulk experiments , we introduce an empirical reduction factor $f$, namely, 1 at $a_w$ = 1, $e^{-1.5}$ at $a_w$ = 0.4 and $e^{-2.5}$ at $a_w$ = 0, and interpolated between $a_w$ = 0 and 0.74 (providing the best agreement with measurement (**Figure 3**)):

$$D_{\mathrm{H_2O,NaCl}}(a_w) = f \times D_{\mathrm{H_2O,NaCl}}(a_w = 0.74)$$

**Equation S2**

Again, $D_{\mathrm{H_2O,NaCl}}(m = 0)$ is given by Zobrist et al.[5]

We approximate the $H_2O$ diffusion coefficient in a ternary aqueous NaCl-sucrose mixture with a sucrose molar ratio of $x = M_{suc}/(M_{suc} + M_{NaCl})$ by a Vignes-type equation:[7,8]

$$D_{\mathrm{H_2O}}(x) = D_{\mathrm{H_2O,\ NaCl}}^{(1-\alpha x)} \times D_{\mathrm{H_2O,\ sucrose}}^{\alpha x}$$

**Equation S3**

With $\alpha = e^{0.8x}$ for $\alpha x < 1$ and $\alpha = 1/x$, for $\alpha x > 1$.

For $Na^+$ and $Cl^-$ ions, in the present study we use the diffusion coefficients as function of water activity and temperature for Synthetic Lung Fluid (SLF) given by Luo et al.[32] This diffusion is independent of sucrose concentration, but this approximation does not prevent good agreement with measurements.

For the diffusivity of $H_2O$ molecules in the gas phase, we use the molecular diffusion coefficient $D_{\mathrm{H2O,gas}}$ = 0.211 cm$^2$/s ($T$/273 K)$^{1.94}$ (1013 hPa/$p$) with temperature $T$ and total air pressure $p$ (Pruppacher and Klett).[9] For the ventilation enhancement factor E, which corrects the molecular diffusion by accounting for eddy diffusion and other air motion, we find that E = 1 yields the best agreement with the experimental data, i.e., no advective enhancement. This is plausible considering that the wells are 10-mm wide and 5 mm deep, which strongly suppresses the formation of eddies within the wells.

#### Model limitations

This study has limitations in three respects: (i) of experimental nature, (ii) in terms of assumptions about the physical processes involved, (iii) approximations made in the biophysical modeling using ResAM.

*(i) Experimental limitations.* The evaporation of the droplets was filmed from a position 15 cm above the well plate. This enables us to precisely evaluate the horizontal projection of the evaporating droplets and the timing of their efflorescence by image recognition techniques. However, it does not allow to measure the flattening of a droplet caused by $H_2O$ loss, when the contact angle between the droplets and the surface of the well plate decreases. Instead, we have to estimate the height of a droplet by calculating its $H_2O$ loss and the resulting mass density. This information enters the modeling with ResAM as outlined in **Figure S4**. Since knowing the solution concentration is of fundamental importance in the context of virus inactivation, future work should aim at improving this by means of a second camera monitoring the droplets from the side.

*(ii) Assumptions about the physical processes involved.* Two fundamental assumptions are made in this work, namely that there is encapsulation of residual water by NaCl crystals in efflorescing pure NaCl solution droplets, and that in NaCl:sucrose droplets microcrystals form that remain too small to be visible. First, the assumption of an encapsulated liquid phase is based on the observational evidence that the inactivation of viruses after efflorescence of droplets exposed to RH = 30% proceeds in agreement with the measured inactivation in a saturated bulk solution (see overlapping red diamond and black dot at 6.1 mol/kg in **Figure 4B**). This virological evidence is also backed up by direct physical evidence using electron microscopy, infrared extinction spectra, and Raman scattering on drying sea salt, sodium chloride and ammonium sulfate aerosol particles.[10–12] This suggests that the formation of liquid pockets that contain saturated solutions is not only possible but likely. Second, the assumption of microcrystal formation in salt solutions with high sucrose content (NaCl:sucrose = 1:8) is necessary to explain why virus inactivation proceeds in accordance with saturated bulk solutions, although the crystals do not grow to visible size. This is probably because the ions cannot travel long distances due to the low diffusivity and, therefore, very many tiny crystals are formed. The resolution of our camera is not much better than 100 µm, which is not sufficient to detect these

crystals. Furthermore, previous studies have shown that organic components can kinetically limit the diffusivity of inorganic ions and the coalescence in liquid-liquid phase-separated particles.[13] in addition, observations in porcine respiratory fluid suggest that the protein matrix is sufficiently viscous to prevent complete coalescence of salts in aqueous solutions.[14] While these studies support the hypothesis of microcrystal formation in the 1:8 NaCl:sucrose solution, it remains a plausible but hypothetical physical process that requires further investigation.

*(iii) Approximations applied in ResAM.* The likely largest modeling uncertainty results from the extrapolations of the concentration dependence of the diffusivities (**Figure 4A**) and inactivation times $t_{99}$ (**Figure 4B**) to highly supersaturated regimes without direct measurements. In pure aqueous NaCl solutions the diffusion coefficients of $H_2O$ molecule are well-constrained by viscosity measurements under subsaturated conditions, i.e., for water activities $a_w > 0.7$ (see the Supporting Text about the Biophysical Modelling for details). In addition, diffusivities have been measured in pure aqueous sucrose for any concentration (see Supporting Text). However, this leaves a considerable degree of freedom for more concentrated NaCl solutions as well as for NaCl-sucrose mixtures. The inactivation rates determined from the three bulk measurements (red diamonds in **Figure 4B**) and the two droplet experiments (black dots) for pure NaCl under subsaturated and mildly supersaturated conditions at $b$ = 0.14, 2.8, 6.1 and 8.0 mol/kg convincingly show the exponential behavior of $t_{99}$ as function of NaCl molality up to a molality of $b \sim 8$ mol/kg. The continuation beyond that concentration with the same exponent is in the first instance an ad-hoc assumption, but as we show in **Figure S9** with the choice of diffusivities shown in **Figure 4A** also the extrapolation of $t_{99}$ to strongly supersaturated conditions up to $b = 20$ mol/kg is very well constrained. However, if the $H_2O$ molecule diffusivity was much higher (and the viscosity much lower), this would in turn lead to a steeper decrease of $t_{99}$ with increasing salt molality, and conversely a diffusivity much lower (and viscosity much higher) would yield a shallower decrease of $t_{99}$ with $b$. A sensitivity test in **Figure S10** shows for strongly supersaturated conditions (water activities $a_w \ll 0.7$) how errors in the diffusivity can be compensated by different slopes in the dependence of $t_{99}$ on the salt molality $b$. The dashed and dotted lines in **Figure S10B** span the range of conceivable uncertainties in of $t_{99}$. Although these uncertainties concern only the strongly supersaturated regime briefly before efflorescence, micron-sized respiratory aerosol particles at typical indoor humidity may reside in this regime for extended times. The uncertainties in $t_{99}$ may then exceed an order of magnitude, calling for improved measurements of the $H_2O$ diffusion coefficient in concentrated salt solutions. – In addition to these uncertainties, other modeling issues appear to be minor. For example, ResAM does not accurately describe the droplet, which is a spherical cap on the bottom of the well and flattens out during the drying process

as the contact angle decreases. Instead, we describe this by a spherical shell, whose thickness corresponds to the time-dependent height of the cap (see **Figure S4**). This procedure allows us to continue using the spherical ResAM without having to create a completely new model. While this approximation works very well near the center of the droplet, the liquid volume is overestimated, and the inactivation is underestimated near the rim of the droplet. This can lead to an overestimation of the titer by a factor of 2 or 3, which is a small error compared to the uncertainties in the $H_2O$ diffusivities.

#### Supporting Figures

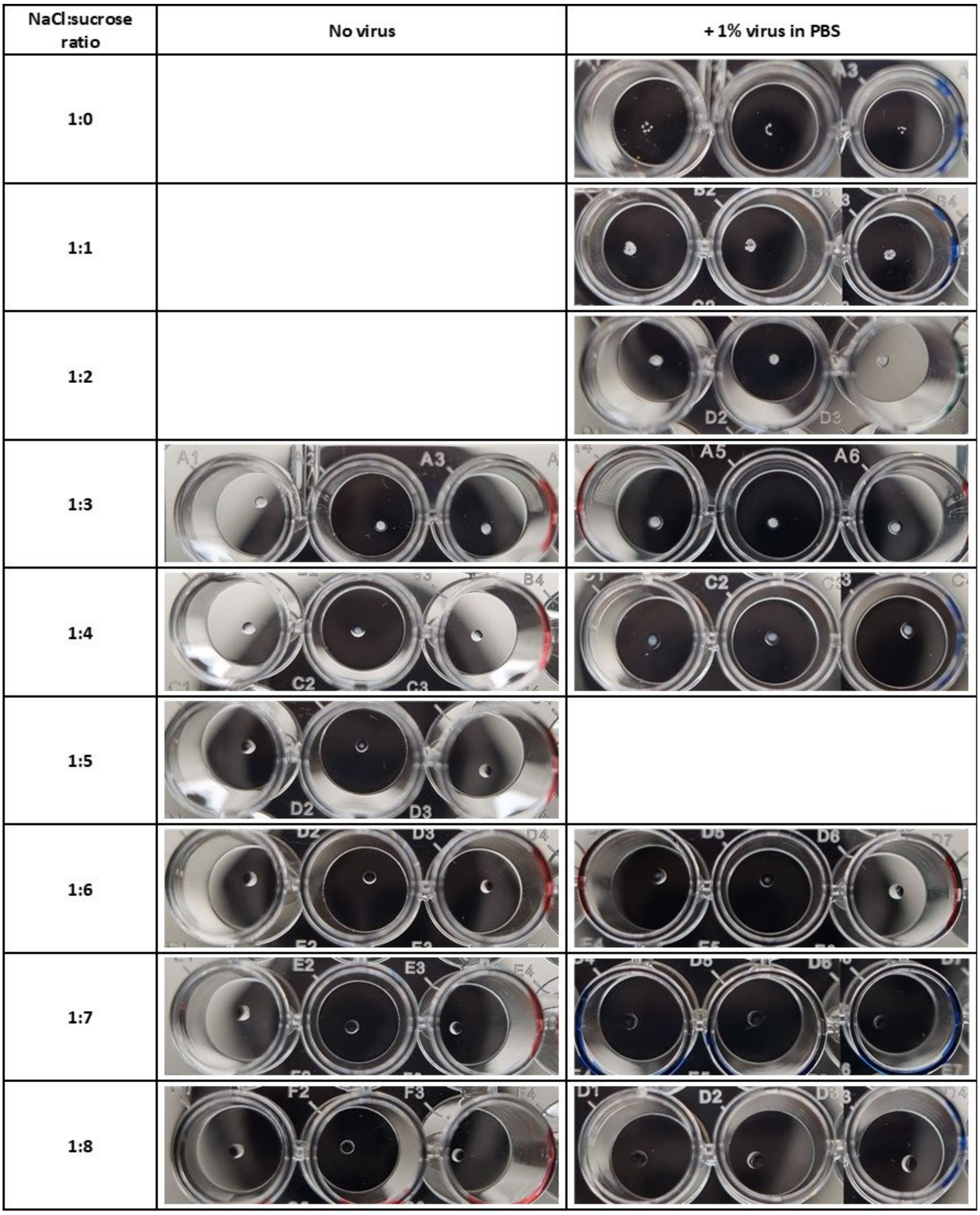


**Figure S1: 1-µl droplets dried for 1 to 3.5 h at 30% RH with various NaCl:sucrose mass ratio**. Various NaCl:sucrose ratios were tested to determine the lowest sucrose content necessary to prevent efflorescence. The initial composition of the droplets was 8 g/l of NaCl and a sucrose concentration according to the indicated NaCl:sucrose mass ratio (i.e., ranging from 0 to 64 g/l). The droplet was either deposited directly ("no virus" column), or spiked prior to deposition with 1% (v/v) of infectious virus in PBS. At ratios 1:0 and 1:1, droplets were fully crystallized. At ratios ranging from 1:2 to 1:5, a crystal core effloresced in the center of the particle and was surrounded by a liquid shell, in both with and without virus conditions. The crystal decreased in size with the increase of sucrose concentration. At ratios ≥1:6, no efflorescence can be seen by eye in the "no virus" condition. A crystal core can be seen in the 1:6 and 1:7 droplets containing viruses. The only ratio not showing efflorescence in the presence of virus was 1:8.

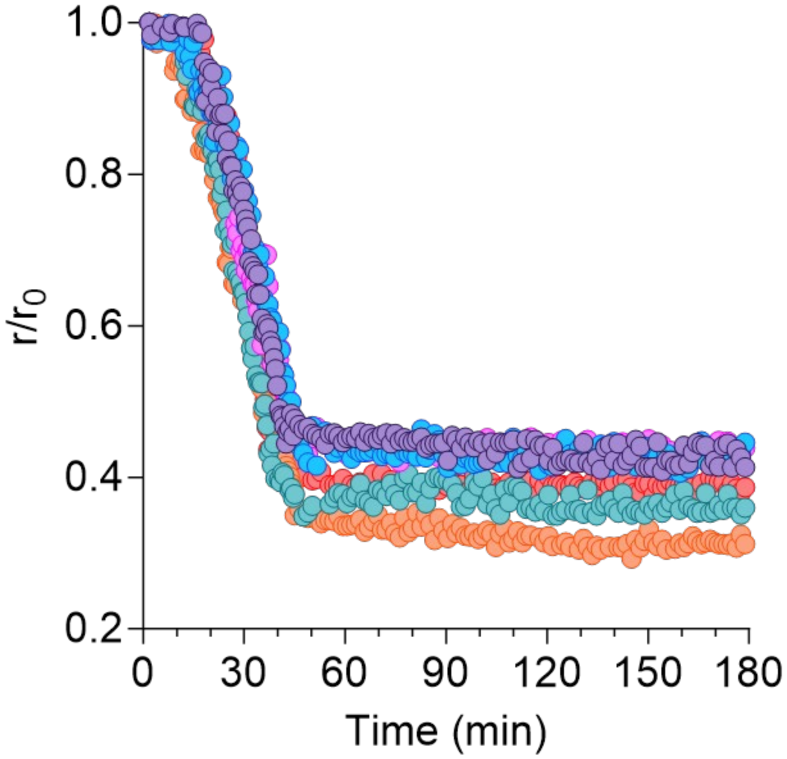


**Figure S2: Normalized NaCl droplet radius shrinkage at 65% RH due to water evaporation until liquid-phase equilibrium with air.** The initial droplet volume was 1 µl and the initial NaCl concentration was 8 g/l. 6 individual droplets were measured simultaneously. Each droplet is displayed in a different color. All droplets reached equilibrium after 45 ± 2 min and remained liquid until the end of the experiment. The analysis of the droplet cap radius has relative standard deviation (coefficient of variation) of 2.4% on average for a single droplet, and of 12% across all droplets. To improve readability, only 1 out of 2 data points are shown in the evaporation phase, and 1 out of 5 data points are shown in the equilibrium phase (tail). The statistics, however, were calculated using all datapoints.

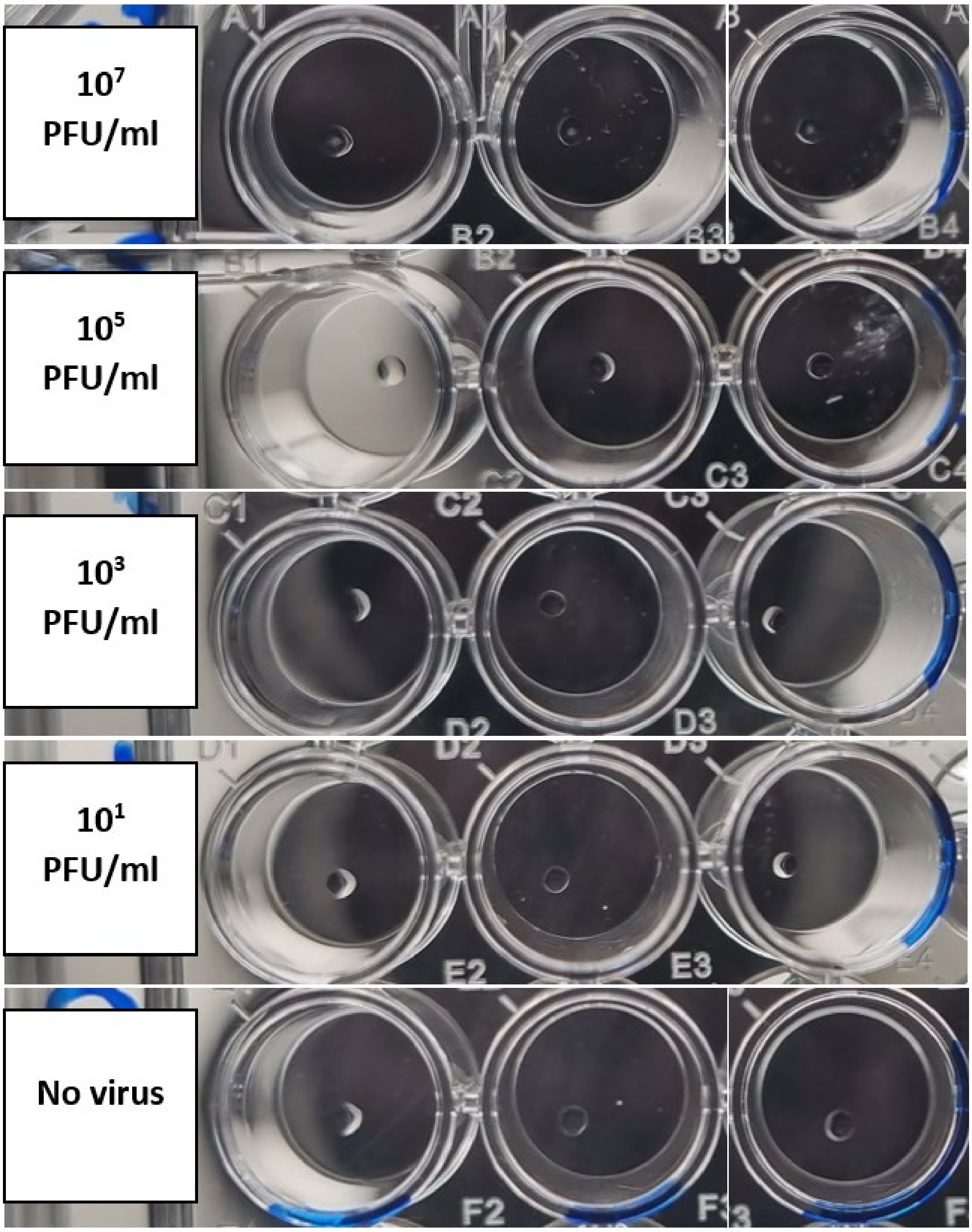


**Figure S3: 1-µl NaCl/sucrose droplets dried for 2 h at 30% RH with various virus titers.** The initial composition of the droplets was 8 g/l of NaCl and 48 g/l of sucrose, reaching a 1:6 NaCl:sucrose mass ratio. The droplets were supplemented prior to deposition with 1% (v/v) of PBS containing between 0 to $10^9$ PFU/ml of viruses, reaching final virus concentrations of 0 to $10^7$ PFU/ml. Efflorescence was observed only in the droplets containing the highest virus concentration ($10^7$ PFU/ml), showing that a minimal amount of virus is required to trigger efflorescence. Control experiments were performed to confirm that the efflorescence was not triggered by the addition of PBS from the viral stock solution (data not shown).

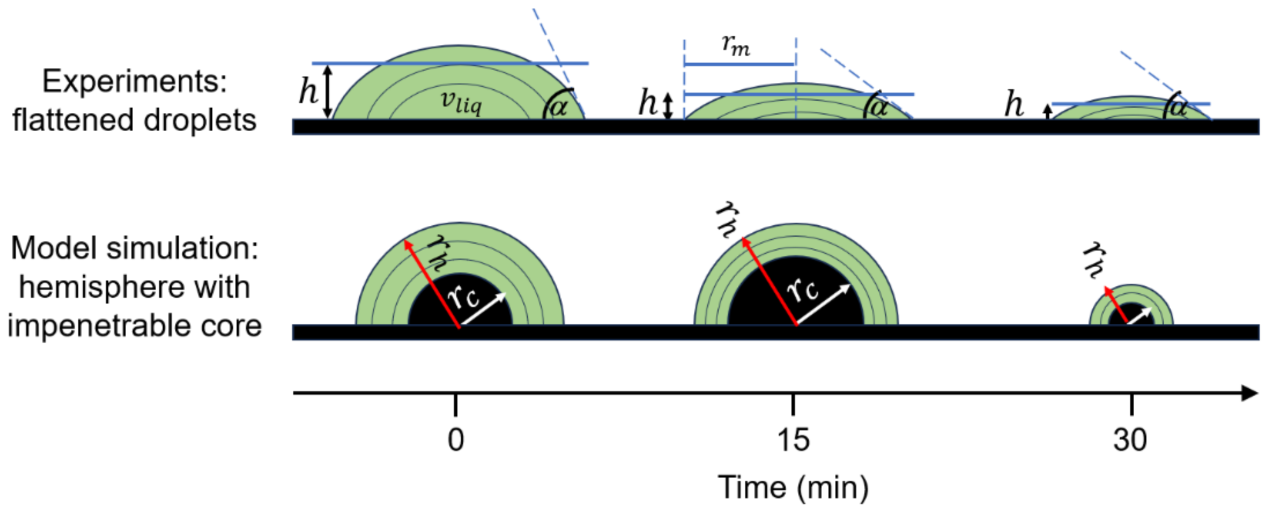


**Figure S4: Modeling of $H_2O$ loss and subsequent IAV inactivation in a droplet with an initial volume of 1 ml deposited on the bottom of the well.** Top row: evaporation of $H_2O$ leading to a flattening of the droplet in the first ~15 min, while the surface area covered by the liquid remains largely unchanged but the contact angle decreases; subsequently, further evaporation leads to shrinkage of the covered surface. Bottom row: model simulation mapping the ellipsoidal droplet shape with average thickness $h$ to a spherical shell of equal thickness on an impenetrable core with radius $r_c$, i.e. $r_h$ - $r_c$ = $h$, which correctly captures the diffusive transport times of $H_2O$ molecules through the liquid matrix (aqueous NaCl solution with varying amounts of sucrose) and provides a good approximation of the solute concentrations. The values of $r_h$, $r_c$ and $h$ are determined from the video images in combination with the liquid volume $v_{\text{liq}} = 2\pi/3\ (r_h^3 - r_c^3)$ calculated by ResAM. ($h$ is calculated analytically from $h = \frac{2}{3r_m^2}\left(r_0^3 - (r_0^2 - r_m^2)^{3/2}\right) - (r_0^2 - r_m^2)^{1/2}$.)

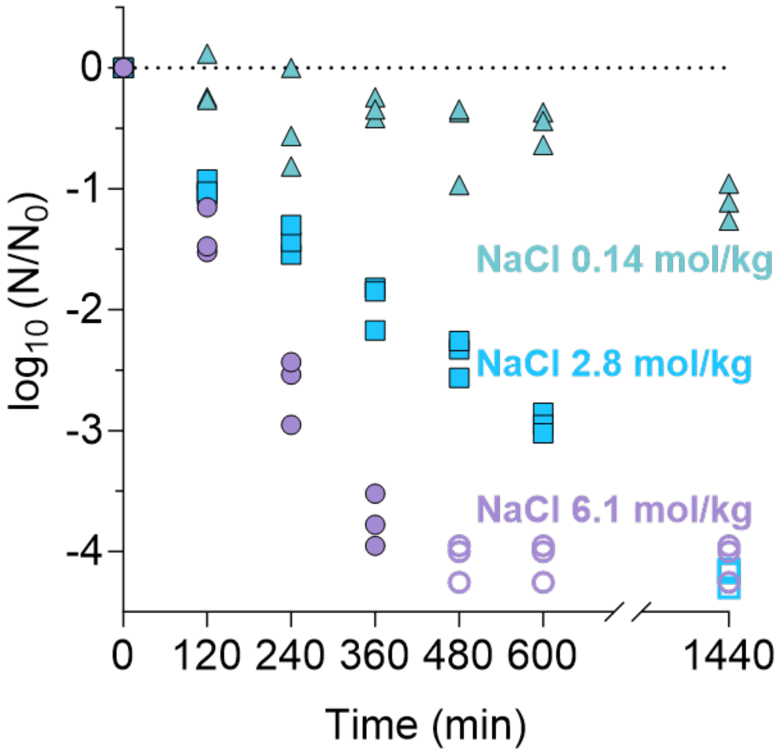


**Figure S5: IAV inactivation in bulk NaCl solutions at 0.14 mol/kg (triangles), 2.8 mol/kg (squares) and 6.1 mol/kg (saturated – circles).** All experiments were conducted in triplicate. Empty symbols indicate data below quantification limit. The corresponding inactivation rate coefficients are listed in **Table S2.** Inactivation can be approximated by first-order kinetics, though close inspection reveals that the decay between the first two time points is slightly more rapid compared to that at later times. This may be due to the presence of a susceptible sub-population that is inactivated rapidly when exposed to inactivating conditions, and is not representative of the overall inactivation rate coefficient. Therefore, this initial decay is neglected and first-order kinetics are assumed for the entire time course.

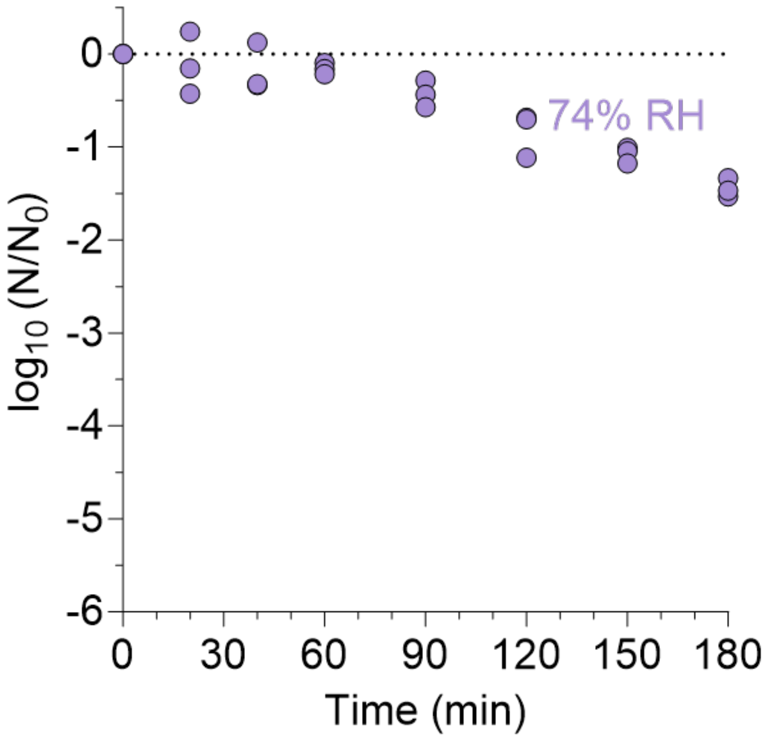


**Figure S6: IAV inactivation in 1-µl NaCl droplets dried at 74% RH.** No efflorescence occurred at this RH. Each dot represents one individual droplet. The initial NaCl concentration was 8 g/l.

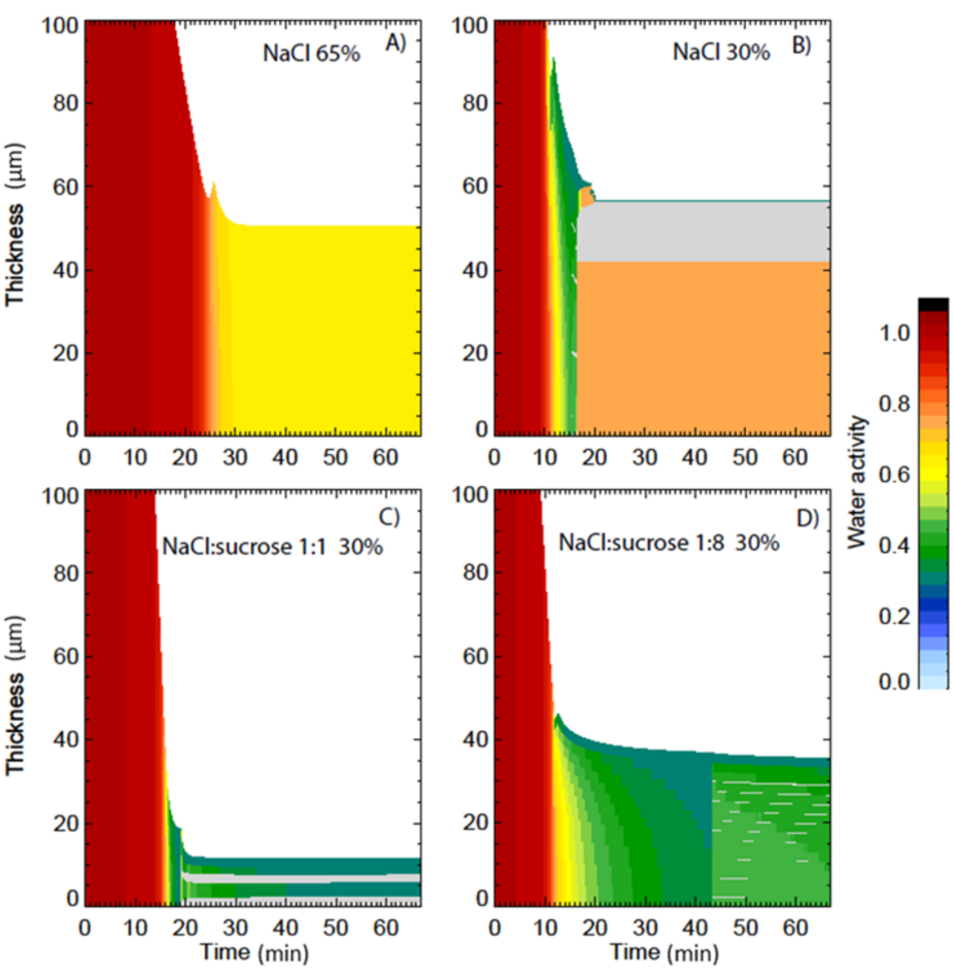


**Figure S7: Evolution of water activity in 1-µl drying droplets, modeled with ResAM.** (A) NaCl droplets at 65% RH; (B) NaCl droplets at 30% RH; (C) NaCl:sucrose 1:1 droplets at 30% RH; (D) NaCl:sucrose 1:8 droplets at 30% RH. Light gray regions show the effloresced NaCl crystals.

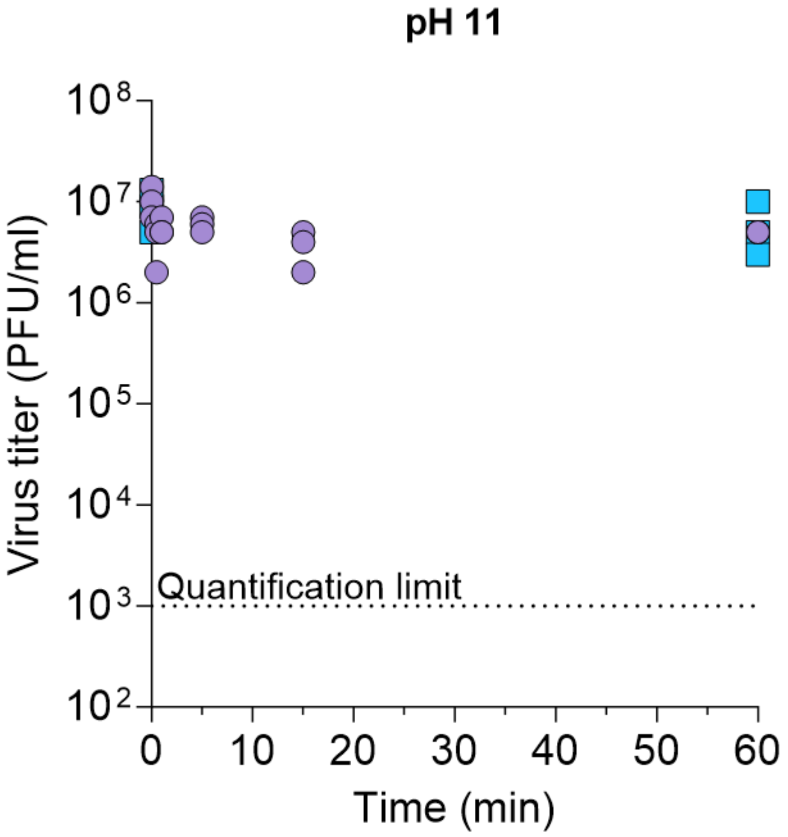


**Figure S8: IAV titer in bulk solution at pH 11.** The solution is composed of 9 ml of trisodium phosphate 0.2 M and 1.5 ml of citric acid 1 M in milli-Q water (measured pH = 11.0 on two different days). All experiments were conducted in triplicate. At each time-point, an aliquot of each sample was diluted 1:100 in PBSi to neutralize the alkaline pH prior to freezing and later titration by plaque assay. Experiments performed on different days are indicated by different symbols.

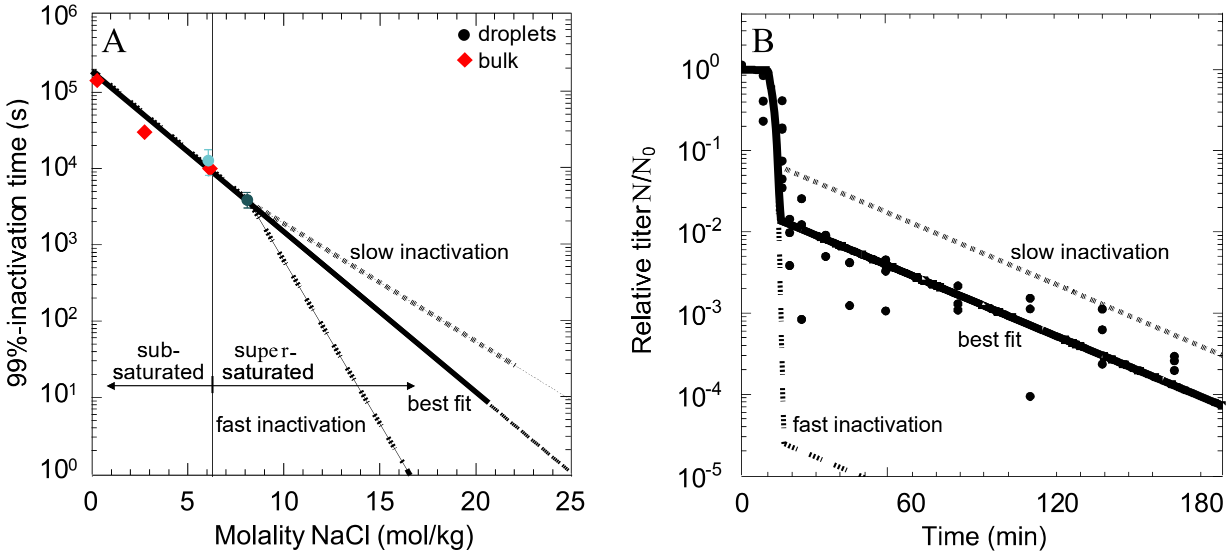


**Figure S9: Sensitivity test for slower (dotted lines) and faster (dashed lines) inactivation in pure aqueous NaCl under strongly supersaturated conditions ($b$ > 8 mol/kg).** The results are compared with the best fit (solid lines) shown in **Figure 4B** and **4H**. (A) $t_{99}$ as function of NaCl molality $b$. (B) Resulting titer during the first 3 hours.

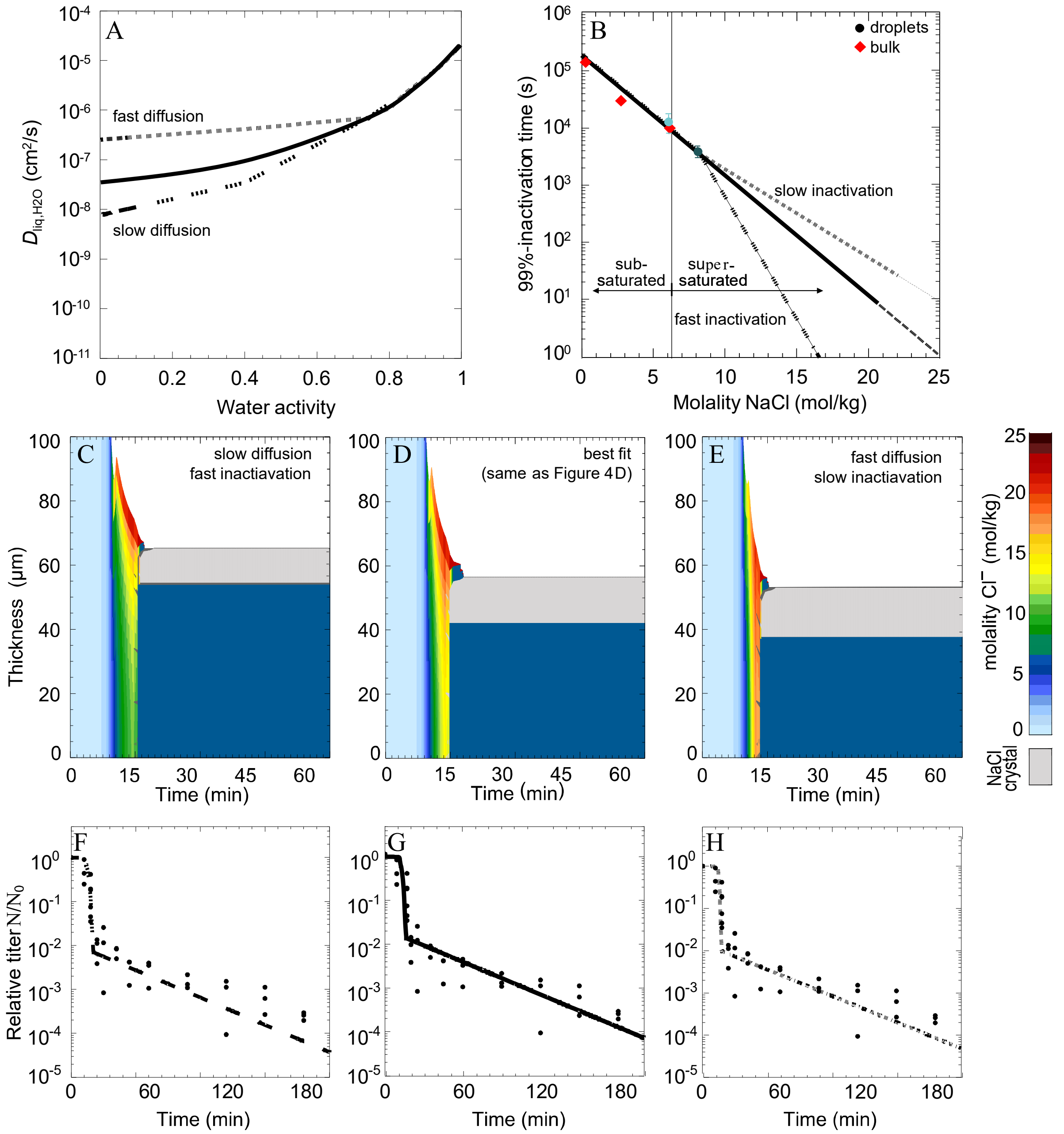


**Figure S10: Sensitivity tests to search for different pairs of $H_2O$ diffusivities and 99%-inactivation times that satisfy the measurements for pure NaCl solutions in the entire sub-and superconcentrated range.** Panels D and G are identical to **Figure 4D** and **G** and identical to the solid curve in **Figure S9B**. Panels C and F show a case with assumed slower diffusion compensated by by faster inactivation, panels E and H the reverse case.

#### Supporting Tables

**Table S1: Check-list of experimental details as requested by MIQE guidelines** according to Bustin et al.[15]

| ITEM TO CHECK | PROVIDED Y/N | COMMENT |
|---|---|---|
| **EXPERIMENTAL DESIGN** | | |
| Definition of experimental and control groups | Y | In materials and methods |
| Number within each group | Y | Specified in figures |
| **SAMPLE** | | |
| Description | Y | In materials and methods |
| Microdissection or macrodissection | NA | |
| Processing procedure | Y | In materials and methods |
| If frozen - how and how quickly? | Y | In materials and methods |
| If fixed - with what, how quickly? | Y | In materials and methods |
| Sample storage conditions and duration (especially for FFPE samples) | Y | In materials and methods |
| **NUCLEIC ACID EXTRACTION** | | |
| Procedure and/or instrumentation | Y | In materials and methods |
| Name of kit and details of any modifications | Y | In materials and methods |
| Details of DNase or RNAse treatment | Y | According to kit instructions |
| Contamination assessment (DNA or RNA) | Y | In materials and methods |
| Nucleic acid quantification | N | Not performed |
| Instrument and method | NA | |
| RNA integrity method/instrument | N | Not performed |
| RIN/RQI or Cq of 3' and 5' transcripts | NA | |
| Inhibition testing (Cq dilutions, spike or other) | Y | In materials and methods |
| **REVERSE TRANSCRIPTION** | | |
| Complete reaction conditions | Y | In materials and methods |
| Amount of RNA and reaction volume | Y | In materials and methods |
| Priming oligonucleotide (if using GSP) and concentration | NA | |
| Reverse transcriptase and concentration | Y | According to kit instructions |
| Temperature and time | Y | In materials and methods |
| **qPCR TARGET INFORMATION** | | |
| Sequence accession number | N | Not provided |
| Amplicon length | Y | In materials and methods |
| *In silico* specificity screen (BLAST, etc) | NA | |
| Location of each primer by exon or intron (if applicable) | NA | |
| What splice variants are targeted? | NA | |
| **qPCR OLIGONUCLEOTIDES** | | |
| Primer sequences | Y | In materials and methods |
| Probe sequences | Y | In materials and methods |
| Location and identity of any modifications | NA | |
| **qPCR PROTOCOL** | | |
| Complete reaction conditions | Y | In materials and methods |
| Reaction volume and amount of cDNA/DNA | Y | In materials and methods |
| Primer, (probe), Mg++ and dNTP concentrations | Y | According to kit instructions |
| Polymerase identity and concentration | Y | According to kit instructions |
| Buffer/kit identity and manufacturer | Y | In materials and methods |
| Additives (SYBR Green I, DMSO, etc.) | Y | According to kit instructions |
| Complete thermocycling parameters | Y | In materials and methods |
| Manufacturer of qPCR instrument | Y | In materials and methods |
| **qPCR VALIDATION** | | |
| Specificity (gel, sequence, melt, or digest) | N | Performed but not reported |
| For SYBR Green I, Cq of the NTC | N | Performed but not reported |
| Standard curves with slope and y-intercept | Y | In materials and methods |
| PCR efficiency calculated from slope | Y | In materials and methods |
| r2 of standard curve | Y | In materials and methods |
| Linear dynamic range | Y | In materials and methods |
| Cq variation at lower limit | Y | In materials and methods |
| Evidence for limit of detection | Y | In materials and methods |
| **DATA ANALYSIS** | | |
| qPCR analysis program (source, version) | Y | In materials and methods |
| Cq method determination | Y | In materials and methods |
| Outlier identification and disposition | NA | |
| Results of NTCs | N | Performed but not reported |
| Justification of number and choice of reference genes | NA | |
| Description of normalisation method | NA | |
| Number and concordance of biological replicates | Y | Specified in figures |
| Number and stage (RT or qPCR) of technical replicates | Y | In materials and methods |
| Repeatability (intra-assay variation) | N | Not performed |
| Statistical methods for result significance | Y | In materials and methods |
| Software (source, version) | Y | In materials and methods |

**Table S2: Inactivation rate coefficients and corresponding 99%-inactivation time in NaCl bulk solution and 1-µl NaCl/sucrose droplets.** The initial NaCl concentration in all droplets was 0.14 mol/kg. For the droplet system, the inactivation rate coefficient corresponds to the inactivation in equilibrated droplets, that is, starting 40 min after deposition. In all systems, the rate coefficient was calculated assuming first order kinetics. Standard errors are indicated.

| **Experimental system** | **Inactivation rate coefficient $k$ ($min^{-1}$)** | **99%-inactivation time $t_{99}$ (min)** | **Corresponding figure** |
|---|---|---|---|
| NaCl bulk solution, 0.14 mol/kg | $1.9 (\pm 0.4)\cdot 10^{-3}$ | $2.4 (\pm 0.5)\cdot 10^{3}$ | **Figure S5** |
| NaCl bulk solution, 2.8 mol/kg | $1.2 (\pm 0.0)\cdot 10^{-2}$ | $3.9 (\pm 0.2)\cdot 10^{2}$ | **Figure S5** |
| NaCl bulk solution, 6.1 mol/kg | $2.5 (\pm 0.1)\cdot 10^{-2}$ | $1.9 (\pm 0.1)\cdot 10^{2}$ | **Figure S5** |
| NaCl droplet, 30% RH | $1.8 (\pm 0.4)\cdot 10^{-2}$ | $2.6 (\pm 0.5)\cdot 10^{2}$ | **Figure 2** |
| 1:1 NaCl:sucrose droplet, 30% RH | $7.6 (\pm 3.3)\cdot 10^{-3}$ | $6.1 (\pm 2.7)\cdot 10^{2}$ | **Figure 3** |
| 1:8 NaCl:sucrose droplet, 30% RH | $6.4 (\pm 1.7)\cdot 10^{-3}$ | $7.2 (\pm 1.9)\cdot 10^{2}$ | **Figure 3** |
| NaCl droplet, 65% RH | $6.5 (\pm 0.4)\cdot 10^{-2}$ | $7.1 (\pm 0.5)\cdot 10^{1}$ | **Figure 2** |
| NaCl droplet, 74% RH | $2.2 (\pm 0.2)\cdot 10^{-2}$ | $2.1 (\pm 0.2)\cdot 10^{2}$ | **Figure S6** |
